# Profiling Allogeneic HLA-specific B-cell Responses Utilizing a 64-plex Single-HLA Reporter Cell Panel

**DOI:** 10.64898/2026.01.19.700453

**Authors:** Shengli Song, Emma F. King, Ashley M. Drabik, Jean Kwun, Cliburn Chan, Annette M. Jackson, Garnett Kelsoe, Stuart J. Knechtle

**Affiliations:** Duke Transplant Center, Duke University School of Medicine, Durham, NC 27710, USA; Department of Surgery, Duke University School of Medicine, Durham, NC 27710, USA; Department of Biostatistics and Bioinformatics, Duke University School of Medicine, Durham, NC 27710, USA; Department of Integrative Immunobiology, Duke University School of Medicine, Durham, NC 27710, USA

## Abstract

Identifying allogeneic HLA-specific B cells in sensitized individuals is essential for defining the cellular basis of allogeneic humoral immunity but remains technically challenging due to their low frequency. To overcome this barrier, we generated a 64-plex single-HLA reporter cell (HLA64-RC) panel that provides a cost-efficient, multiplex, high-throughput platform for screening B-cell specificity. We additionally developed a companion R package, *HLA64*, for automated data analysis and visualization. Integrated with a streamlined high-throughput BCR discovery workflow, this platform enables reliable identification and characterization of allogeneic HLA-specific B cells from sensitized transplant candidates. In a pilot application, thirteen HLA-specific B cells were identified, enabling linked analyses of phenotype, function, and BCR genetics. These B cells exhibited an IgG^+^ CD24^low^ phenotype, diverse HLA allele-specificity profiles, and recurrent heavy- and light-chain V-gene usage. In two independent B-cell lineages, clonal members within each lineage displayed divergent binding patterns despite sharing a common clonal origin. Broader application of this approach for systematic profiling of alloreactive B-cell responses will help elucidate the molecular basis of allorecognition, define immunodominant HLA eplets, and ultimately improve immunological risk assessment and allograft outcomes in transplant recipients.

## INTRODUCTION

Humoral responses are involved in different types of allograft rejection [1], and antibody-mediated rejection (AMR) is the most common immunological cause of graft failure in kidney transplantation [2–4]. Pre-existing or *de novo* donor-specific antibodies (DSAs) against the mismatched allogeneic HLA molecules are an independent risk factor for AMR and poor long-term allograft outcome [5, 6]. The use of sensitive immune assays that detect anti-HLA antibodies (Abs) in serum has substantially improved pre- and post-transplant alloimmune risk stratification [7]. However, these assays capture only the terminally differentiated plasma cell (PC) compartment and do not assess memory B cells (MBCs), a long-lived population capable of rapid expansion and differentiation into antibody-secreting cells (ASCs) upon antigen re-encounter [8].

Over the past two decades, two major types of methods have been developed to profile allogeneic HLA-specific B cells. The first uses soluble fluorescent HLA probes to directly detect antigen-specific B cells. This intuitively straightforward approach, however, suffers from high background labeling and low specificity for truly antigen-specific B-cell receptors (BCRs) [9–11].

The second type of approach infers MBC specificity after *in vitro* polyclonal activation, followed by detection of secreted Abs using ELISpot [12, 13], FluoroSpot [14], or clinical HLA single-antigen bead (SAB) assays [15]. These assays revealed that HLA-specific MBCs can persist for decades [16–18] and may be detectable even when the corresponding serum Abs are absent [11, 15, 17–19]. Moreover, work from the Bestard group associated donor-specific MBC frequencies with AMR severity [17, 19] and identified donor-reactive MBC expansion as an independent predictor of AMR in non-sensitized patients [19]. Because these assays measure secreted Abs rather than B cells directly, they exhibit high analytical sensitivity and specificity. However, they provide no information on the phenotype or BCR genetics of individual B cells, and their resolution is limited, as ELISpot and FluoroSpot measure only a few specificities at once and SAB data from polyclonal cultures may be difficult to interpret when multiple reactivities are detected.

In parallel with these efforts in B-cell profiling, remarkable progress has been achieved in HLA molecular mismatch analysis to improve immunological risk stratification in clinical donor–recipient pairing. Instead of considering HLA matching based on the whole HLA antigen, the term “eplet” was used to define small polymorphic amino acid motifs within 3.0-3.5 Å on the HLA molecular surface, representing potential structural determinants recognized by BCRs or Abs [20]. Many studies have demonstrated the association between HLA eplet mismatch load and *de novo* DSA formation [21–24]. Employing matching strategies based on HLA eplets rather than HLA antigens in solid organ transplantation may also increase the donor pool and improve transplant outcome for highly HLA-sensitized patients [25, 26]. However, these eplets are computationally defined, with only a small portion being validated by allo-reactive monoclonal Abs (mAbs) [27]. Determining the immunogenicity and antigenicity of individual eplets or epitopes is therefore essential before eplet-based matching can be adopted clinically [28–30].

A critical step toward such validation is the isolation of individual allo-reactive BCRs from sensitized individuals. This need has renewed interest in direct HLA-probe–based B-cell identification. Using fluorescent HLA monomer, tetramer or dextramer probes, a series of studies from the Heidt and Kramer group have identified five DR-specific mAbs from three [31], 15 DQ-specific mAbs from five [32], and two class I-specific and seven class II-specific mAbs from four sensitized individuals [33], collectively confirming over ten eplets listed in the HLA Eplet Registry [34]. A recent study from the Lund group used dual-color A*01:01 tetramers to isolate 49 mAbs from the graft and peripheral blood of a patient experiencing AMR, enabling detailed structural characterization of epitope recognition [35]. These studies demonstrate the feasibility of probe-based strategies for isolating allo-reactive BCRs and advancing our understanding of the immunogenicity of allogeneic HLA epitopes.

Despite these successes, isolating truly HLA-specific B cells remains inefficient. Early studies reported high background binding [9–11], and even with improved probes, only a minority of probe-binding cells represent truly antigen-specific B cells. For example, in one study [31], 40–60 million peripheral blood mononuclear cells (PBMCs) were processed per subject, yet only 3.4–30.6% of DR-tetramer–positive B cells proved to be genuinely specific. Given the high polymorphism of HLA molecules and the diversity of sensitization histories, large-scale, systematic profiling of allo-reactive BCR repertoires will require high-throughput strategies capable of screening thousands of single-B-cell cultures or cloned mAbs in a multiplex manner. Although clinical SAB assays are highly multiplexed, sensitive, and standardized, their high per-sample cost limits their feasibility for high-throughput screening (HTS) applications in basic research. In addition, false-positive binding to denatured HLA antigens presented on bead surfaces [36–40] remains a concern in research settings. Thus, a key bottleneck in allo-reactive BCR discovery is the absence of a cost-efficient, scalable assay that measures binding to native HLA molecules in a format compatible with multiplex, single-B-cell–based screening.

Here, we present such a solution: a cost-efficient, multiplex single-HLA binding assay designed for HTS applications that utilizes native HLA molecules displayed on the surface of mammalian cells. Building on our previously established multiplex reporter cell platform [41, 42], we generated a 64-plex single-HLA reporter cell panel that provides broad coverage of common HLA alleles across diverse U.S. populations. We confirmed cell surface expression of corresponding HLA molecules and their specificity in Ab detection. We also developed a companion R package for automated data analysis and visualization to further facilitate HTS applications. Finally, by integrating this assay with a recently optimized BCR discovery platform [43], we successfully identified allogeneic HLA-specific B cells from sensitized transplant candidates. The linked phenotypic, functional, and BCR genetic analyses of these cells provide a foundation for deeper investigation into the mechanisms and determinants of allogeneic HLA-specific B-cell responses.

## RESULTS

### Selection of representative HLA alleles for reporter cell panel development

To design a generalizable reporter cell panel for systematic profiling of allogeneic HLA-specific B cells, a key consideration is selecting a limited yet representative set of HLA alleles from among the tens of thousands of allelic variants identified across global populations. Our goal was to maximize both population-level HLA allele coverage and representation of the theoretical HLA eplet repertoire, which underlies Ab specificity at the structural level. To achieve this, we selected the most frequent HLA-A, -B, and -DRB1 alleles across five genetically diverse U.S. subpopulations (Supplementary Fig. 1) using data from the Allele Frequency Net Database [44] (Supplementary Data 1). For DRB3/4/5, DQA1 and DQB1, allele frequency data from a collection of global populations [45] were used (Supplementary Data 1) due to limited U.S.-specific datasets. Additionally, alleles were chosen to ensure broad representation across HLA gene families and cross-reactive epitope groups (CREGs) [46].

Based on these criteria, we designed a 64-plex allele panel (Fig. 1a). Compared with the clinical SAB panel, which achieves 74.3–99.9% population coverage at the allele level and 99.8–100% coverage at the HLA gene family level, the 64-plex panel provides 46.7–96.5% allele-level coverage and 75.2–100% gene family-level coverage across diverse populations (Fig. 1b, c). Despite the reduced number of alleles, the 64-plex panel captures the majority of the theoretical eplet repertoire (Supplementary Data 2): 200/229 (87.3%) HLA-A/B eplets, 120/124 (96.8%) HLA-DRB eplets, and 81/83 (97.6%) HLA-DQ eplets (Fig. 1d). For comparison, the clinical SAB panel covers 226/229 (98.7%) HLA-A/B, 124/124 (100%) HLA-DRB, and 82/83 (98.8%) HLA-DQ eplets. Thus, the 64-plex panel achieves a high level of immunogenetic coverage while dramatically reducing assay cost (approximately 1/500 of the SAB assay), making it suitable for mechanistic and discovery-based research applications.

**Fig. 1.**
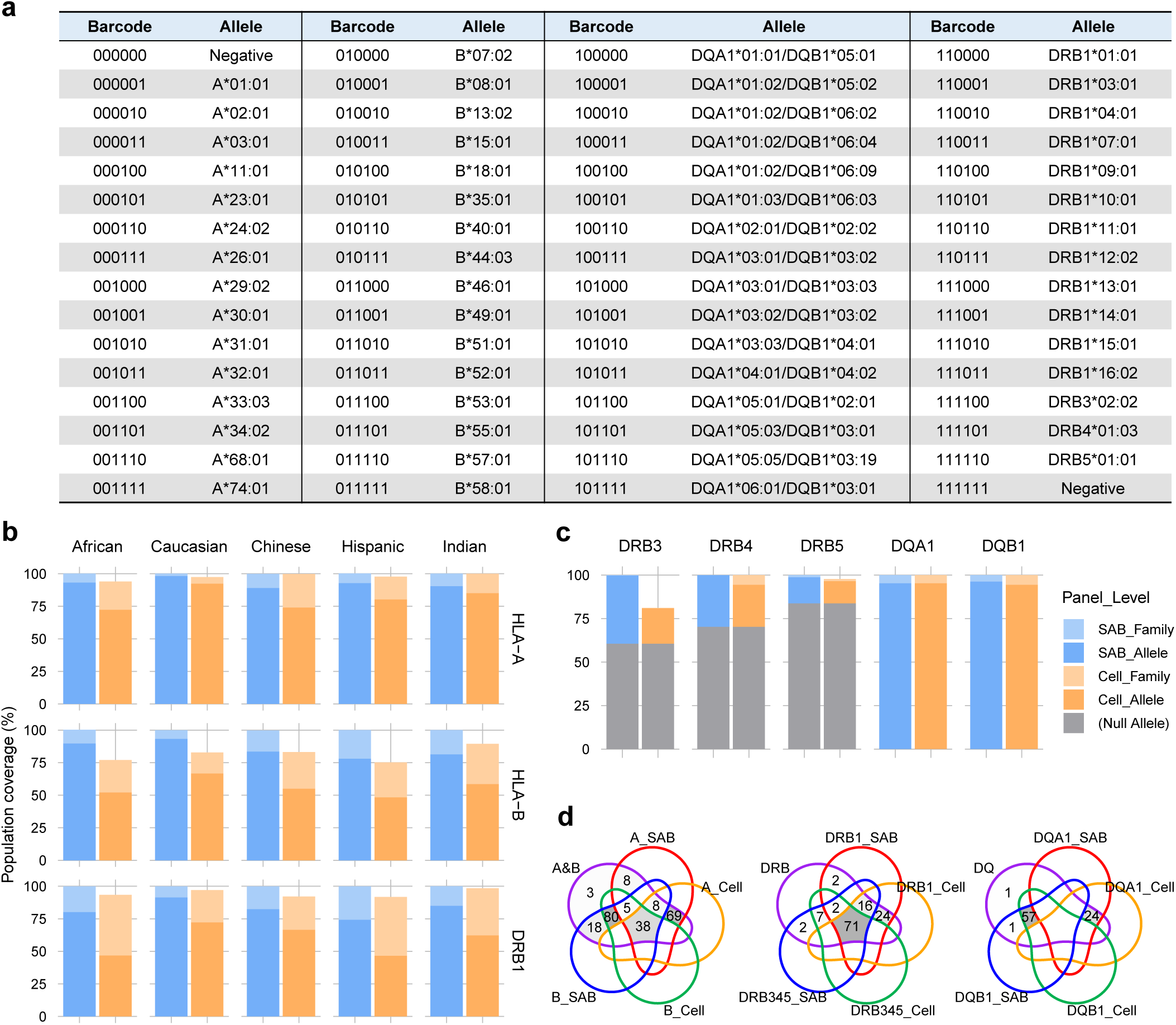
Selection of representative HLA alleles for reporter cell panel development. a) Selected HLA alleles and their corresponding FP barcodes used for the generation of the reporter cell panel, including 15 HLA-A, 16 HLA-B, 16 HLA-DQ, and 15 HLA-DR alleles, along with two negative control cell lines expressing none or all six FPs but lacking HLA expression. b) Comparison of population coverage for HLA-A, -B, and -DRB1 loci between the clinical SAB panel (blue) and the designed 64-plex reporter cell panel (orange). Percentages of coverage at the allele level (dark) and gene-family level (light) across five genetically diverse U.S. subpopulations (Supplementary Fig. 1 and Supplementary Data 1) are shown. c) Population coverage for DRB3, DRB4, DRB5, DQA1, and DQB1 loci by the SAB and designed reporter cell panels, shown as in (**b**). Allele frequency data from a collection of global populations were used (Supplementary Data 1). d) Representation of the theoretical HLA eplet repertoire (Supplementary Data 2) captured by the SAB and designed reporter cell panels for combined HLA-A and -B (left), combined DRB1 and DRB3/4/5 (middle), and combined DQA1 and DQB1 (right) loci. Individual regions in the Venn diagrams are labeled with the numbers of eplets in each compartment and shaded in grey gradients according to their proportions among the total number of eplets for the corresponding combined loci.

### Generation and demultiplexing of the 64-plex single-HLA reporter cell (HLA64-RC) panel

We extended the previously developed cell-based multiplex binding assay platform [41, 42] from 16-plex to 64-plex by careful selection of six fluorescent proteins (FPs) with distinct emission spectra as combinational barcodes, including EBFP2 [47], mTurquoise2 [48], LSSmOrange [49], hmKeima8.5 [50], NowGFP [51], and mKelly1 [52]. Fig. 2a illustrates the experimental process for generating the HLA64-RC panel expressing the selected HLA alleles. First, the HLA-negative K530 cell line, a *CD32A*-depleted derivative of K562 cells [41], was repeatedly transduced with different combinations of the six FPs mentioned above, termed FP barcodes. Sixty-four (2^6^ = 64) basal reporter cell lines were established, each expressing a distinct FP barcode. Monoclonal lines were generated by single-cell sorting and selected to ensure uniform and stable expression of individual FPs. Subsequently, each basal reporter cell line was transduced with a unique HLA allele, resulting in an HLA64-RC panel corresponding to the 64-plex HLA allele panel described above. Finally, these individually barcoded cell lines were pooled at equal ratios and used in multiplex binding assays, followed by detection by multicolor flow cytometry and demultiplexing during data analysis. Fig. 2b shows an example of demultiplexing flow cytometry data obtained with the established HLA64-RC panel. Following successive gating on the six FP channels, 64 subpopulations were successfully identified with unambiguous resolution.

**Fig. 2.**
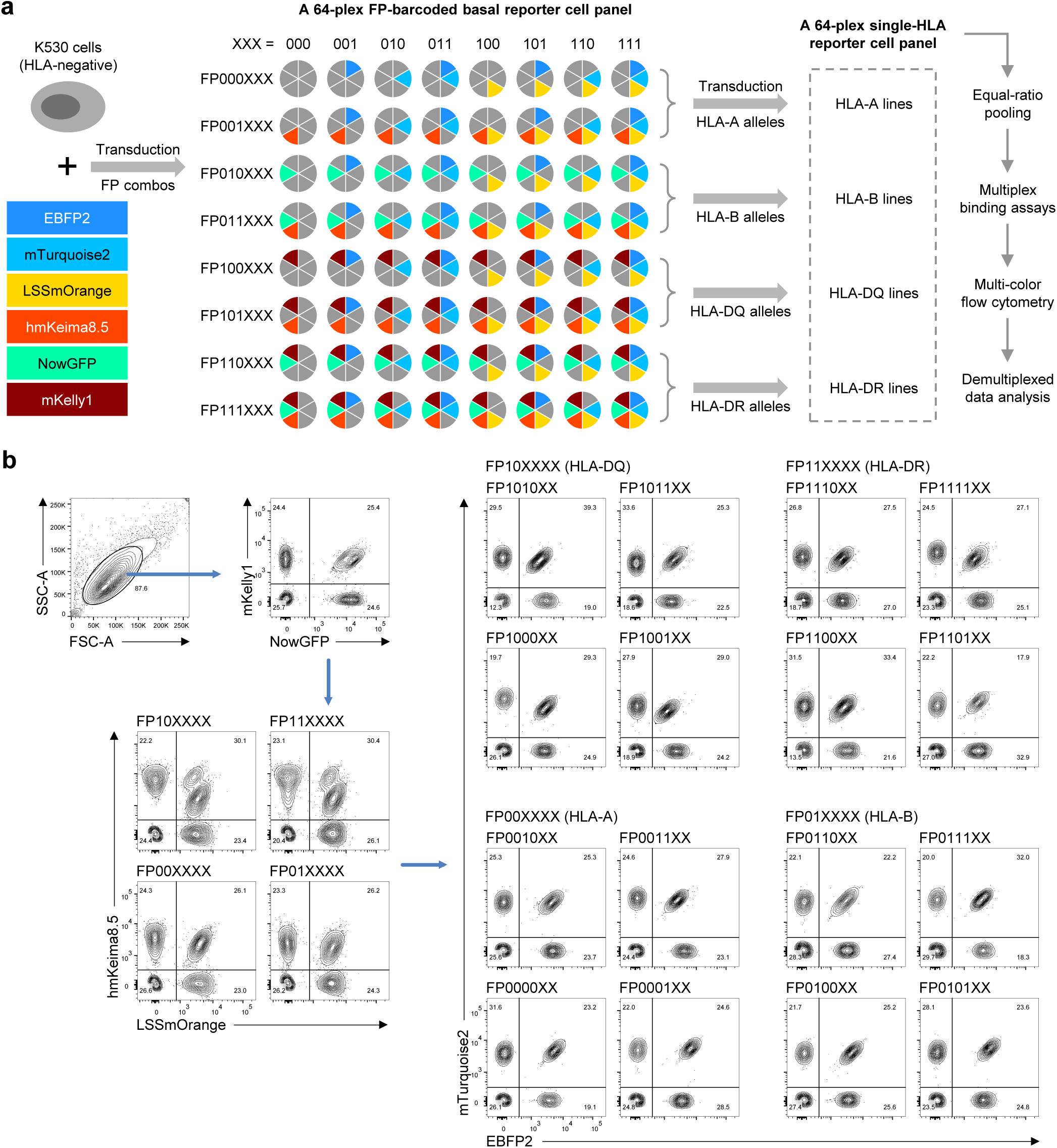
Generation and demultiplexing of the HLA64-RC panel. a) Schematic overview of the experimental process for generating the HLA64-RC panel. The pie charts represent K530-based basal reporter cell lines expressing distinct combinations of six FPs, color-coded according to their peak emission wavelengths. b) Representative example showing the demultiplexing of flow cytometry data acquired using the HLA64-RC panel by sequential gating on the six FP channels.

### Confirmation of single-HLA molecule expression on demultiplexed reporter cell lines

With the established HLA64-RC panel, we performed multiplexed binding assays using reference mAbs with defined pan- or allele-specific reactivities to confirm the surface expression of corresponding HLA molecules on individual cell lines after demultiplexing. As shown in Fig. 3, labeling with human IgG1 κ and λ isotype control Abs resulted in minimal background after secondary detection with PE-conjugated polyclonal anti-human IgG, confirming assay specificity. In contrast, the humanized mAb BB7.2 specifically bound to the A02:01-expressing reporter cell line, demonstrating the utility of this platform for detecting HLA-specific Abs in analyses of allogeneic humoral responses. Additional HLA class I–reactive reference Abs tested included the pan-class I W6/32, Bw4-specific REA274, Bw6-specific REA143, A03:01-specific GAP.A3, and B*07:02-specific BB7.1, all of which demonstrated reactivity to their corresponding allele-expressing cell lines. The slightly elevated background labeling by W6/32 across most class II–transduced lines is likely due to low-level expression of endogenous HLA-C and HLA-E molecules, as previously reported for the parental K562 cells [53]. This weak pan-reactivity can be readily distinguished during analysis of allogeneic responses involving selected HLA alleles.

**Fig. 3.**
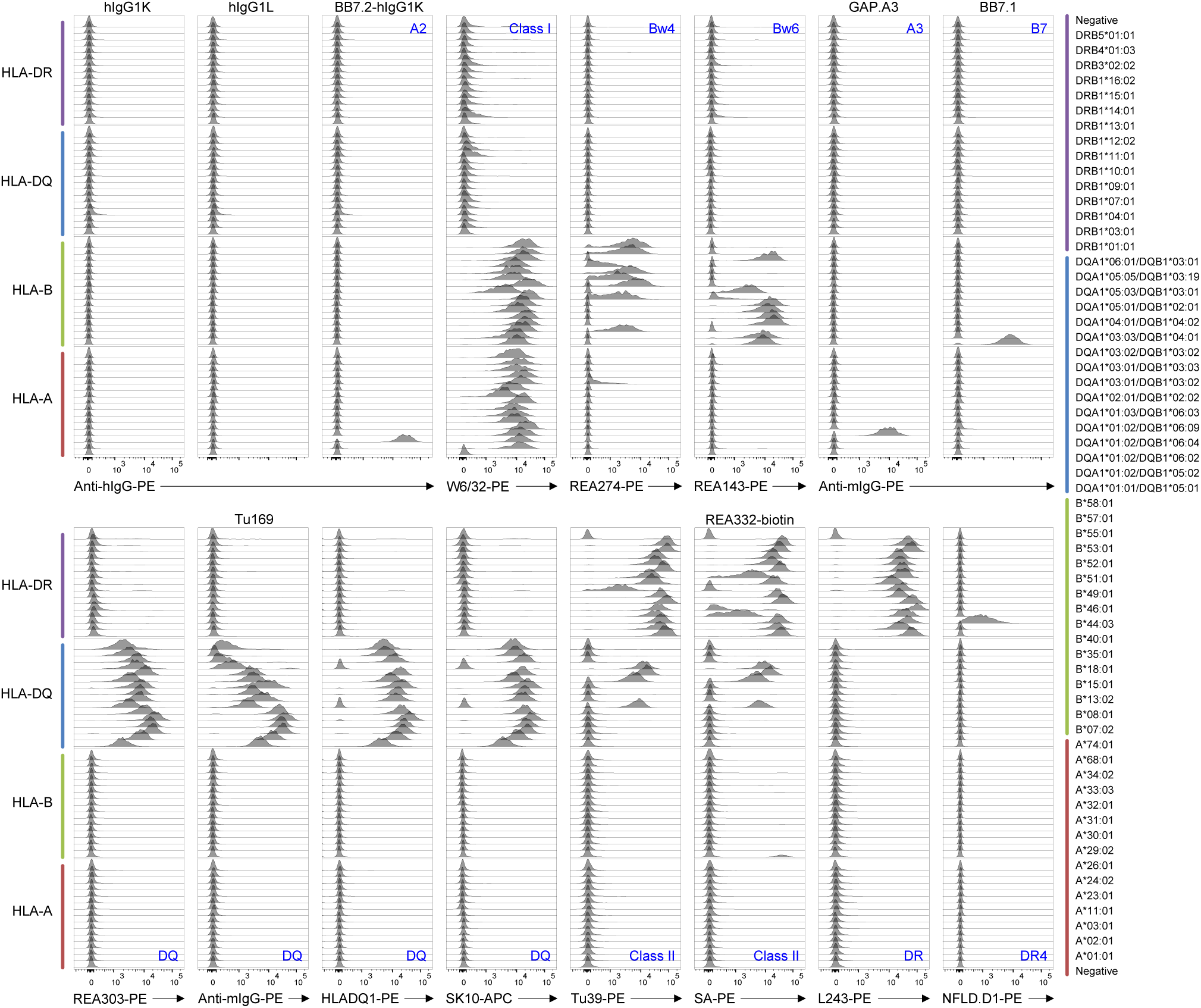
Confirmation of single-HLA molecule expression on demultiplexed reporter cell lines. The HLA64-RC panel was labeled with fluorochrome-conjugated Abs or with unconjugated primary Abs followed by fluorochrome-conjugated secondary Abs. Following flow cytometry acquisition, data were analyzed using *FlowJo*™ software. Live cell events were demultiplexed into 64 populations based on the six FP channels as in Fig. 2b. Stacked histograms of the demultiplexed populations show fluorescence intensities corresponding to Ab labeling. Histograms are grouped by HLA-A (dark red), HLA-B (green), HLA-DQ (blue), and HLA-DR (purple) alleles, displayed upward in the order indicated on the far-right. Blue labels positioned in the upper-right (top row) or lower-right (bottom row) corners denote the reported specificities of the corresponding mAbs. hIgG1K and hIgG1L, human IgG1κ and IgG1λ isotype control Abs. BB7.2-hIgG1K, BB7.2 in human IgG1κ format. Anti-hIgG-PE and anti-mIgG-PE, PE-conjugated anti-human or anti-mouse IgG secondary Abs. SA-PE, PE-conjugated streptavidin.

For HLA-DQ and -DR expressing lines, the pan-DQ-specific Ab REA303 recognized all DQ lines, whereas Tu169 showed reduced binding to certain DQ7 and DQ2 alleles. HLADQ1 and SK10 displayed identical reactivity patterns, both lacking recognition of the two DQ2 alleles included in the panel, consistent with previous reports [54]. In contrast, the pan-class II Abs Tu39 and REA332 reacted with the same DQ2 and DQ4 alleles but exhibited distinct DR reactivity profiles (Fig. 3). The pan-DR-specific Ab L243 bound strongly to all DR alleles across the panel, while the DR4-specific Ab NFLD.D1 reacted exclusively with DRB1*04:01. Together, these results confirm cell-surface expression of the transduced HLA class II alleles on the reporter cell lines.

### Comparison of the specificities of HLA64-RC and SAB assays using serum samples

To further assess the specificity of the HLA64-RC assay, we analyzed ten serum samples from sensitized transplant candidates whose clinical SAB reactivities covered all HLA alleles included in the HLA64-RC panel. Serial dilutions of serum samples were prepared, and the HLA64-RC assays were performed blinded to the clinical SAB data. IgG binding data at all dilutions are shown in Supplementary Document 1, and Supplementary Fig. 2 presents representative samples at dilutions yielding the highest signal-to-background ratios.

Median fluorescence intensity (MFI) values were compared between the HLA64-RC (Supplementary Data 3) and SAB (Supplementary Data 4) assays for shared alleles (Fig. 4a). Among 327 SAB-detected reactivities, 200 (61.2%) were captured by the HLA64-RC assay. As aggregated in Fig. 4b, the log_10_(MFI) values showed strong correlations between the two assays, with Lin’s concordance correlation coefficients (CCC) ranging from 0.75 to 0.93 and Spearman correlation coefficients from 0.77 to 0.94. The reactivities undetected by the HLA64-RC assay were predominantly those with low MFI values in the SAB assay, as reflected by the shallow slopes in regions with low SAB MFI values (Fig. 4b). Consistently, standardized major axis (SMA) regression yielded slopes of 0.76–1.02 across different HLA loci. Together, these results indicate that the HLA64-RC and SAB assays exhibit concordant specificities, although the HLA64-RC assay shows moderately reduced sensitivity.

**Fig. 4.**
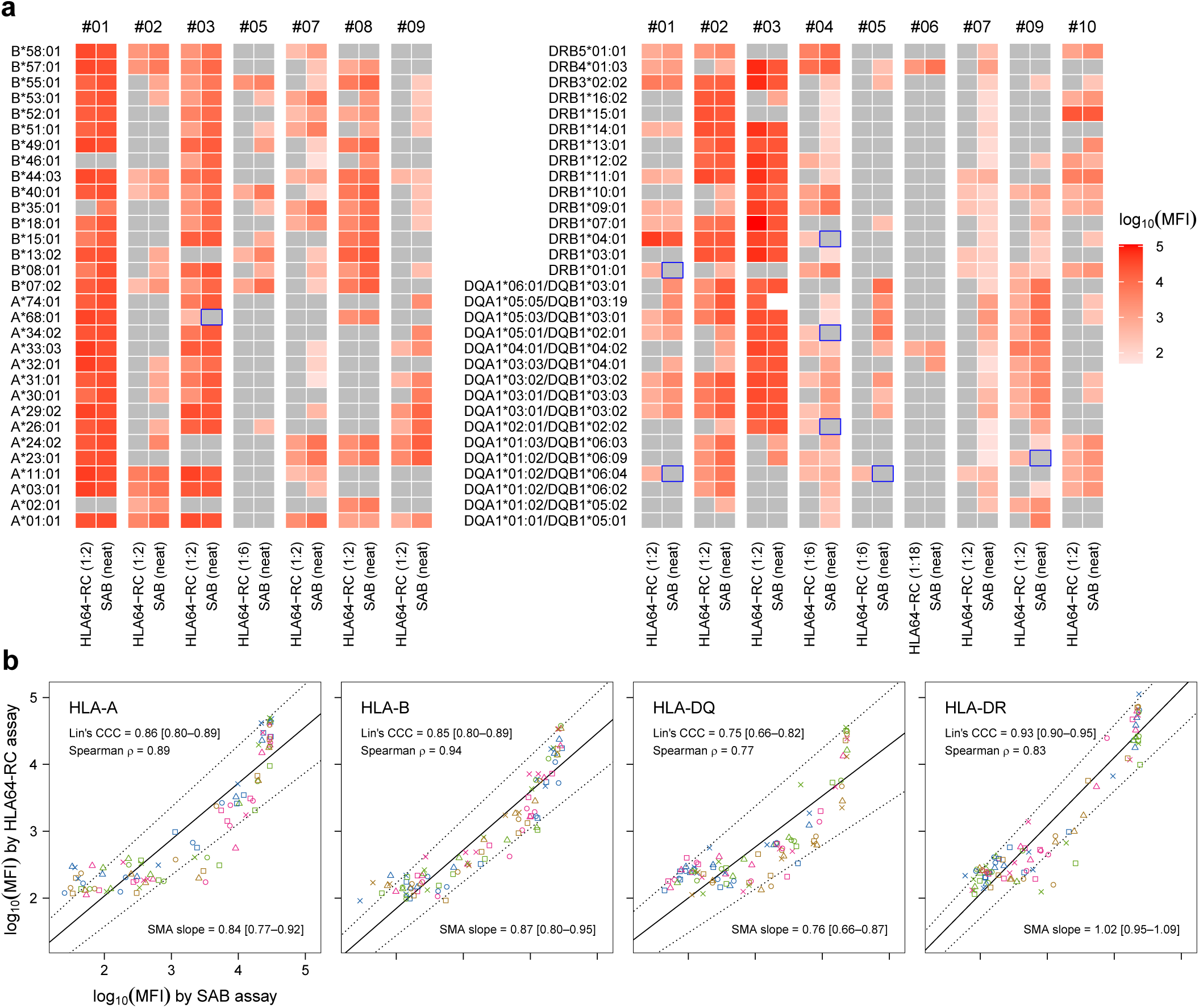
Comparison of the specificities of HLA64-RC and SAB assays using serum samples. Using the HLA64-RC panel, ten serum samples (#01–#10) from sensitized transplant candidates were tested for HLA-binding activities at 1/3 serial dilutions from 1:2 to 1:486, followed by secondary labeling using PE-conjugated anti-human IgG at 2 µg/ml. Raw binding data are shown in Supplementary Document 1. The dilutions yielding the highest signal-to-background ratio for individual samples were selected (Supplementary Fig. 2) and MFI values (Supplementary Data 3) were compared with those obtained from clinical SAB assays (Supplementary Data 4) using the same samples. a) Heatmaps comparing log_10_(MFI) values for shared alleles between the HLA64-RC and SAB assays. The selected dilutions for the HLA64-RC assay were indicated in parentheses. For the SAB assay, serum samples were tested neat. Alleles with MFI values below the binding threshold (see Methods) are shown in grey. Reactivities detected by the HLA64-RC assay but absent in the SAB assay are highlighted with blue boxes. b) Dot plots comparing log_10_(MFI) values for shared alleles between the HLA64-RC and SAB assays, as in (**a**). Data from multiple samples were pooled and plotted separately for HLA-A, -B, -DQ, and -DR loci, with color and shape indicating individual alleles within each locus. For consistency, only serum samples tested at 1:2 dilution in the HLA64-RC assay in (**a**) were included. Within each locus, Lin’s CCC with 95% bootstrap confidence interval (CI), Spearman correlation, and SMA regression with 95% bootstrap CI were calculated using log_10_(MFI) values from the two assays. Solid diagonal lines represent SMA regression fits; dotted lines indicate the 95% CI.

Several non-exclusive factors may contribute to the seemingly lower sensitivity of the HLA64-RC assay (also see Discussion). First, serum samples were diluted at least 1:2 to reduce background binding at higher concentrations (Supplementary Document 1), which would diminish detection of low-affinity Abs. Second, antigen density on the cell surface may be lower than on bead surfaces, particularly for DQ loci [55], consistent with the lowest correlation coefficients and SMA slope for DQ (Fig. 4b). Third, denatured HLA antigens on bead surfaces [36–40] can produce false-positive signals in SAB assays. Notably, eight reactivities detected by the HLA64-RC assay were absent in SAB assays (highlighted in blue in Fig. 4a), emphasizing structural or conformational differences between bead-immobilized recombinant HLA and native, membrane-anchored HLA molecules.

### Development of the *HLA64* R package for automated data analysis and visualization

With the HLA64-RC panel established, we applied this assay in HTS experiments in mechanistic studies for an ongoing clinical trial. One major challenge we experienced was the processing, visualization, and identification of HLA allele binding patterns with the flow cytometry data derived from high-volume and multiplexed sample testing. We therefore developed an R package, *HLA64*, with a user-friendly graphic interface for automated data analysis and visualization to further facilitate HTS applications.

Supplementary Fig. 3 illustrates the overall workflow using the *HLA64* R package. The data processing procedures were visually verified using example flow cytometry data from Fig. 2b. Briefly, the raw flow cytometry data were imported, and live-cell gating was set up automatically (Fig. 5a). After logicle transformation [56, 57], cutoff values were determined automatically for each FP channel to separate negative and positive populations (Fig. 5b), allowing unambiguous demultiplexing of 64 cell populations (Fig. 5c). We also verified the procedures for demultiplexed HLA-specific Ab binding data visualization using example flow cytometry data from Fig. 3. The binding patterns exported from the *HLA64* R package (Fig. 5d) were consistent with those exported from the FlowJo™ software (Fig. 3). Furthermore, re-analysis of the serum binding data from Fig. 4 demonstrated that the MFI values of demultiplexed 64 populations across ten samples were highly concordant with those analyzed using the FlowJo™ software, with Pearson and Spearman correlation coefficients of 1.00 across different HLA loci, and slopes ranging from 1.00 to 1.01 after linear regression fitting (Fig. 5e).

**Fig. 5.**
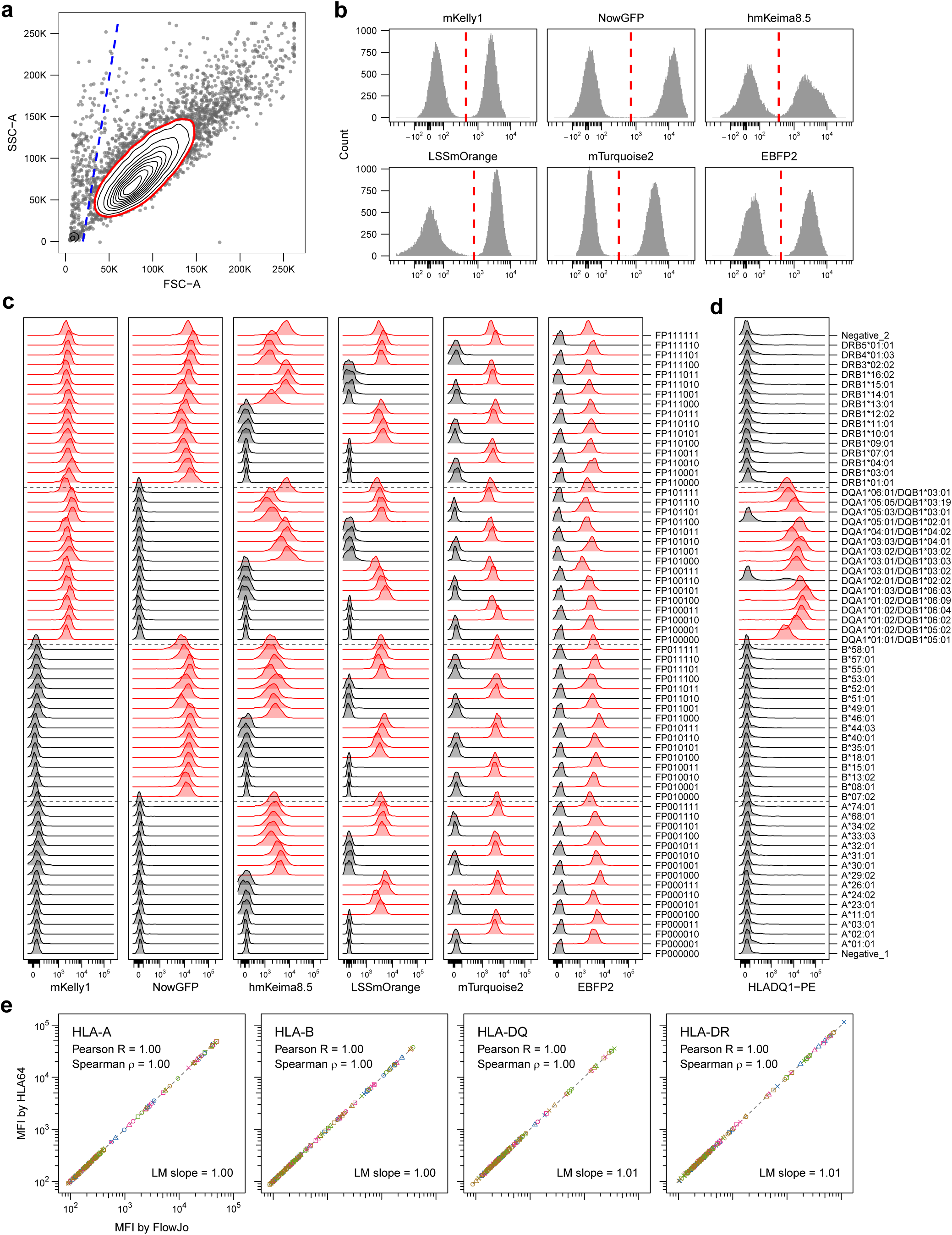
Development of the *HLA64* R package for automated data analysis and visualization. The *HLA64* R package was developed to enable automated analysis of data generated from the HLA64-RC assay. Example flow cytometry data from Fig. 2b were used for side-by-side comparison in panels (a–c). a) Automated live-cell gating (red polygon) based on a two-dimensional kernel density estimate (KDE) of FSC and SSC signals. The blue dashed line indicates a user-defined cutoff to exclude debris and dead cells. b) Automated determination of cutoff values (red dashed lines) for six FP channels by fitting a two-component Gaussian mixture model to distinguish FP-expressing and non-expressing populations. c) Verification of FP expression patterns across the six FP channels for the 64 automatically demultiplexed populations. FP barcodes assigned to individual populations are shown on the far right. Populations with fluorescence intensities above the cutoffs established in (**b**) are highlighted in red. d) Example of binding data output analyzed using the *HLA64* R package. The same flow cytometry data analyzed with *FlowJo*™ in Fig. 3 were used for comparison. Demultiplexed HLA alleles corresponding to individual populations are indicated on the right. Populations exceeding the binding threshold (see Methods) as compared with an unlabeled control sample are highlighted in red automatically. Dashed lines separate HLA loci in the panel. e) Comparison of MFI values obtained using *FlowJo*™ and the *HLA64* R package from the same patient serum binding data shown in Fig. 4. Data from all ten samples were pooled and plotted separately for HLA-A, -B, - DQ, and -DR loci, with color and shape indicating individual alleles within each locus. For each locus, Pearson and Spearman correlations and linear model (LM) regression analyses were performed based on MFI values from both analyses. Dashed diagonal lines represent LM regression fits.

When we applied the *HLA64* R package to data analysis in the HTS experiments mentioned earlier using the HLA64-RC assay, the processing of flow cytometry data from twenty 96-well plates took only a few minutes using the parallelized multi-batch processing mode from the package on a regular desktop PC equipped with 24 logical processors. In comparison, this process typically takes about five hours when using the FlowJo™ software. Summarizing binding patterns across those twenty plates took another few minutes. Depending on the complexity of the binding patterns, the same analysis based on FlowJo™ exports typically takes from half an hour to several hours of manual processing, which can limit throughput and reproducibility. Therefore, we have established the *HLA64* R package, which enables high-throughput and automated data analysis and visualization, yielding highly concordant outputs with improved time efficiency compared with the process using the FlowJo™ software. The user-guided automated workflow also contributes to reproducibility of the analysis.

### Identification and characterization of allogeneic HLA-specific B cells from sensitized transplant candidates

We then applied the HLA64-RC assay and automated data analysis tool in HTS experiments to identify and characterize allogeneic HLA-specific B cells from sensitized transplant candidates. Briefly, patient samples were selected based on the HLA allele reactivities obtained from serum SAB tests. B cells enriched from patient-derived PBMC samples were labeled with corresponding soluble HLA monomer antigen probes, together with fluorochrome-conjugated B-cell phenotyping Abs. Monomer-binding B cells were index-sorted by flow cytometry into single B-cell cultures using a newly optimized method [43] that builds upon a previously established approach [58] to promote differentiation to ASCs. HLA-binding B-cell cultures were identified by screening immunoglobin (Ig)-positive culture supernatants in high-throughput HLA64-RC assays. HLA-specific BCRs were cloned from identified B-cell cultures, and recombinant Abs (rAbs) were produced for further characterization.

Fig. 6 shows representative results from one patient sample in a pilot experiment. Using two phenotyping Ab panels in parallel, B cells were gated as CD19^+^ IgD^−^ monomer^+^ for single B-cell sorting and culture (Fig. 6a). A permissive gating was applied to include monomer-binding B cells with weak fluorescence signals, based on our previous experience. From 823 single B-cell cultures, 445 Ig^+^ cultures were recovered, yielding a cloning efficiency of 54.1%. Using the HLA64-RC assay, seven HLA-binding cultures were identified from 270 Ig^+^ Panel 1 cultures, and six from 175 Ig^+^ Panel 2 cultures (Fig. 6b and Supplementary Fig. 4). The low positivity rate (2.6–3.4%) of truly HLA-specific B cells among monomer-binding B cells highlights the limited specificity of these HLA probes in binding to specific BCRs. The weak binding signals observed (Fig. 6b) necessitated the use of permissive gating to maximize recovery, albeit at the cost of further reducing positivity and efficiency.

**Fig. 6.**
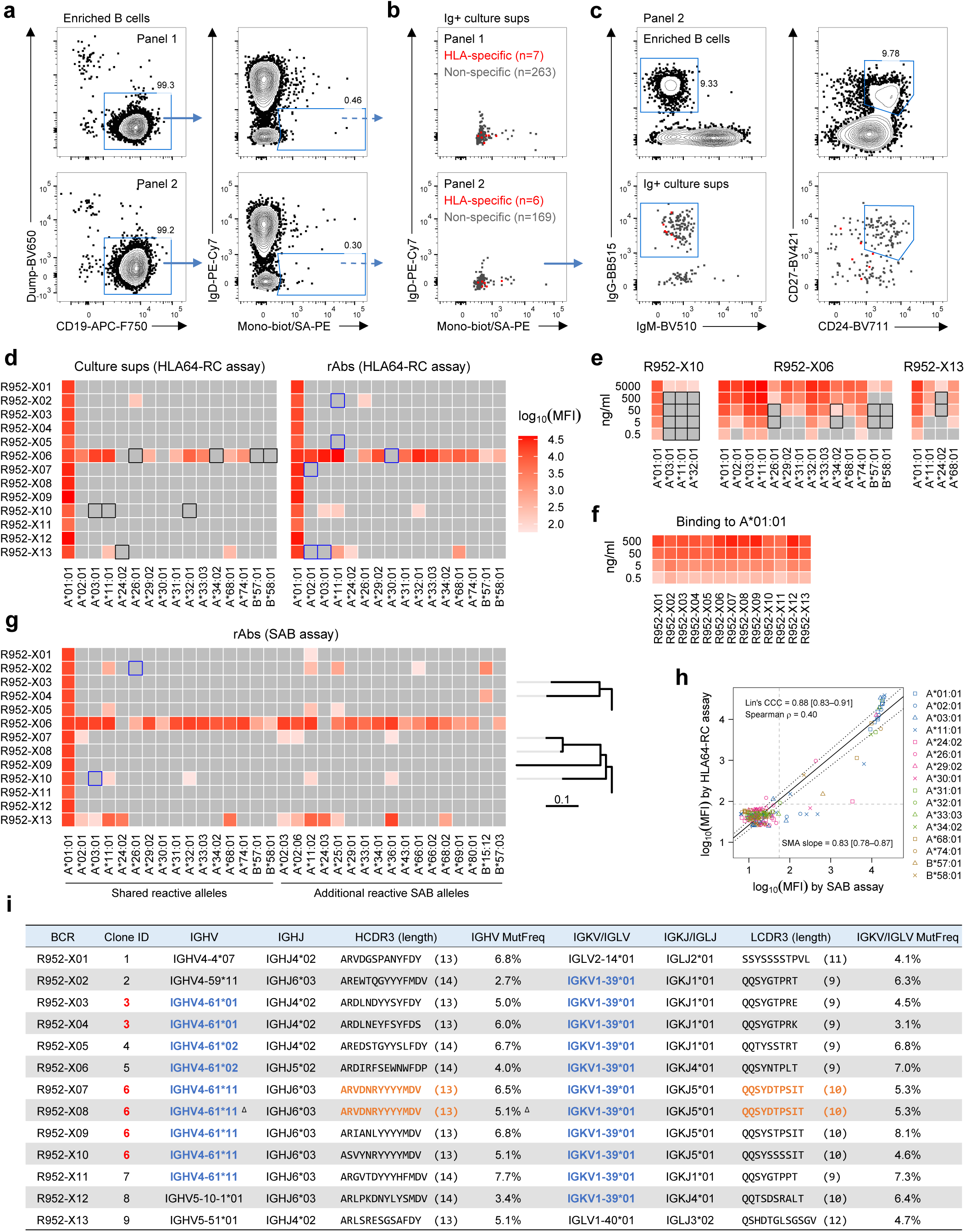
Identification and characterization of allogeneic HLA-specific B cells from sensitized transplant candidates. a) Gating strategy for identifying HLA monomer-binding B cells for single-cell sorting and culture. Two phenotyping Ab panels were tested in parallel (see Methods). b) HLA monomer-binding intensities of index-sorted, monomer-binding single B-cell cultures that were Ig^+^ in the supernatants. HLA-specific cultures are highlighted in red. c) Phenotypic analysis with additional labeling Abs in Panel 2. Top row shows enriched total B cells; bottom row shows index-sorted, monomer-binding Ig^+^ single B-cell cultures as in (**b**), with HLA-specific in red. d) Heatmaps show log_10_(MFI) HLA-binding values of identified HLA-specific single B-cell cultures (left) and corresponding rAbs (5 µg/ml; right) analyzed by HLA64-RC assays. Only reactive alleles are shown; those below binding thresholds (see Methods) are in grey. Black boxes in culture group mark reactivities undetected in cultures but present in rAbs; blue boxes in rAb group mark reactivities undetected in HLA64-RC assays but present in SAB data in (**g**). e) Serial dilutions of rAbs corresponding to the three cultures with undetected reactivities in (**d**) recapitulated the same binding patterns (black boxes). f) Heatmaps show log_10_(MFI) binding values to the major reactive allele A*01:01 for all 13 rAbs at serial dilutions. g) Heatmaps show log_10_(MFI) binding values of rAbs (5 µg/ml) tested by SAB assays. Shared or additional reactive alleles are indicated. Blue boxes denote reactivities that were undetected as compared with the rAb group in (**d**). Phylogenetic trees (right) align corresponding clonal members of two identified B-cell lineages shown in (**i**). h) Comparison of log_10_(MFI) values for shared reactive alleles between HLA64-RC and SAB assays using rAbs, as in (**d**) and (**g**). Lin’s CCC, Spearman correlation, and SMA regression were calculated. The solid line shows SMA regression fit; dotted lines indicate 95% CI; dashed lines mark binding cutoffs. i) V(D)J and clonal analyses of the 13 HLA-specific Abs. Clone IDs of the two B-cell lineages are in red; shared IGHV/IGLV usages in blue; identical HCDR3/LCDR3 sequences in orange; Δ denotes replacement with a lower-score IGHV call matching clonal members.

Based on the index-sorting data, we retrospectively examined the B-cell phenotypes using the additional markers included in Panel 2. All six identified HLA-specific B cells were IgG^+^ class-switched; however, none displayed a CD24^hi^ CD27^+^ typical memory phenotype (Fig. 6c). Additional phenotyping markers will be required to better define these cells. Analysis of additional patient samples will be necessary to determine whether this phenotype is reproducible and to clarify its biological relevance.

BCR sequences were cloned from the thirteen HLA-specific B-cell cultures, and rAbs were produced. HLA64-RC assays performed using these rAbs (Supplementary Document 2) yielded binding patterns concordant with those obtained from the corresponding culture supernatants during the initial screening (Fig. 6d and Supplementary Data 5). The eight undetected reactivities (highlighted in black) from three cultures were likely due to lower Ab concentrations in the supernatants than the purified rAbs used, as serial dilutions of corresponding rAbs recapitulated the same patterns, with the eight reactivities becoming undetectable at lower concentration ranges (Fig. 6e). These results demonstrate that the combination of the single B-cell culture method and the HLA64-RC assay provides an effective and reliable approach for identifying HLA-specific B cells, even with HLA probes of limited specificity.

These rAbs allowed us to evaluate the sensitivity and specificity of the HLA64-RC assay at the mAb level. As shown in Fig. 6f, for the major reactive allele A*01:01, the HLA64-RC assay was capable of detecting binding activities at concentrations as low as 0.5 ng/ml for all thirteen rAbs tested, demonstrating good sensitivity for a cell-based binding assay. We also compared the binding patterns of these rAbs tested in the HLA64-RC (Supplementary Data 5) and clinical SAB (Supplementary Data 6) assays at identical concentrations. Concordant binding patterns were obtained across shared alleles (Fig. 6d, g), with a Lin’s CCC of 0.88 between log_10_(MFI) values from the two assays (Fig. 6h). These assays also revealed substantial diversity in epitope recognition within this small set of rAbs, with nine distinct binding patterns classified by the SAB assay among the thirteen rAbs (Fig. 6g).

Similar to the case in serum sample testing (Fig. 4), six SAB reactivities were undetected by the HLA64-RC assay, and two reporter cell reactivities were absent in the SAB results (highlighted in blue in Fig. 6d, g). Unlike the case with polyclonal sera, the use of mAbs allowed us to dissect this discordance at the individual allele level. For instance, the SAB reactivity to A*11:01 was undetected by two rAbs (R952-X02 and -X05) but was detected by another rAb (R952-X10) in the HLA64-RC assay, even though the SAB MFI value of this reporter-positive rAb was lower than those of the two rAbs that failed to show reactivity in the HLA64-RC assay. Another example involved allele A03:01, where two rAbs showed opposing patterns: R952-X13 was detected only by the SAB assay, whereas R952-X10 was detected only by the HLA64-RC assay. These observations further support the notion that structural or conformational differences exist between recombinant HLA molecules immobilized on bead surfaces and native membrane-anchored HLA molecules on mammalian cell surfaces. Such differences may underlie the moderate SMA slope and reduced Spearman correlation observed between assays (Fig. 6h).

V(D)J clonal analysis of these thirteen HLA-specific BCRs provided several interesting observations (Fig. 6i). First, among nine clonotypes (seven singletons and two multi-member lineages) identified, five shared a common IGHV gene segment (IGHV4-61), and seven shared an IGKV allele (IGKV1-39*01), although the IGHJ and IGKJ gene usage and CDR3 lengths varied. This recurrent V-gene usage suggests that germline-encoded binding determinants within Ig V gene segments, analogous to the germline-encoded amino acid–binding (GRAB) motifs described in immunodominant public antiviral epitope responses [59], may also contribute to allogeneic HLA recognition. Second, all BCRs underwent somatic hypermutation in their heavy and light chain V gene segments, with IGHV mutation frequencies ranging from 2.7–7.7% and IGKV/IGLV mutation frequencies ranging from 3.1–8.1%, consistent with germinal center (GC)-experienced B cells circulating in the periphery. Third, combined analysis of phylogenetic trees for the two B-cell lineages and their corresponding SAB binding patterns (Fig. 6g) revealed that clonal members within the same lineage exhibited distinct allele-binding profiles. This finding suggests limitations in the current approach of predicting HLA eplets based on allele binding patterns and highlights the possible contribution of polymorphic amino acid residues outside core HLA eplets in mediating allogeneic recognition and shaping humoral responses [35, 60]. Lastly, the two members (R952-X07 and -X08) of clone 6 shared identical heavy and light chain CDR3 sequences yet displayed distinct binding patterns, consistent with the role of V gene-encoded residues in shaping HLA recognition.

## DISCUSSION

We designed, developed, and validated the HLA64-RC assay as an alternative to the clinical SAB assay for basic research applications, including profiling allogeneic humoral responses and identifying or characterizing HLA-specific BCRs and Abs. The HLA64-RC assay offers multiple distinct advantages. (1) Native antigenicity. HLAs are displayed as full-length, membrane-anchored molecules on a human-derived cell line, preserving their native conformation and antigenicity as present on primary human cells. (2) Generalizability. The panel was intentionally constructed to provide broad coverage of common HLA alleles and gene families across diverse populations, with strong representation of predicted eplet repertoires across loci, making it widely applicable for studies of allogeneic humoral immunity. (3) Expandability. Multiplex detection by multicolor flow cytometry permits straightforward expansion: additional HLA alleles can be incorporated along with compatible FPs [41] or by labeling sub-panels with cell-reactive dyes (e.g., CellBrite® NIR680) prior to pooling. Multiple Ig classes or subclasses can also be measured in parallel using differentially conjugated secondary Abs [61]. Specifically expanded panels covering unrepresented HLA epitopes in ethnic populations [62] could serve as a supplement to mainstream SAB assays in clinical applications. (4) Accessibility. Because the assay is built on standard flow cytometry rather than Luminex^®^ platforms, it is more accessible to most research laboratories. (5) Convenience. Individual reporter cell lines can be expanded, pooled at defined ratios [41], and cryopreserved as stock vials, enabling batch-consistent, ready-to-use reagents. (6) Cost-efficiency. Production and expansion of stock vials require minimal resources, and the estimated per-test cost is ∼1/500 that of the clinical SAB assay. There is potential for this assay, after further development and standardization, to function as an alternative to SAB assays, particularly for patients in low-income countries. (7) HTS-amenability. The low assay cost, combined with high-throughput sampling capabilities on modern flow cytometers and a companion automated data analysis and visualization pipeline (the *HLA64* R package), supports scalable HTS applications.

One limitation of the HLA64-RC panel is its more limited HLA allele coverage compared with SAB panels, such that reactivities directed against alleles not represented in the panel will be undetected. Consequently, in its current format, the panel is intended for research use only and is constrained by its predefined allele composition. Importantly, this platform is inherently modular, and targeted expansion of the panel is feasible, allowing additional HLA alleles to be incorporated as needed to tailor the panel for specific research or clinical applications.

Another limitation of the HLA64-RC assay is its seemingly moderately reduced sensitivity compared with the clinical SAB assay (Figs. 4 and 6). This may reflect lower antigen density on the cell surface, resulting in a higher detection threshold in terms of Ab concentration or binding affinity. From another perspective, however, the artificially high HLA density on SAB beads can exaggerate Ab binding and generate false-positive or clinically less-relevant signals [55, 63]. Denatured HLA molecules and cryptic neo-epitopes on bead surfaces [36–40] may further contribute to false-positive signals in SAB assays [63], explaining some reactivities undetectable in the HLA64-RC assay. Importantly, the ability of the HLA64-RC assay to detect reactivities absent in SAB assays (Figs. 4a and 6g) argues against a purely sensitivity-based explanation and instead supports the presence of structural or conformational differences between bead-immobilized and cell-surface-expressed HLA molecules. When applied to serum testing (Fig. 4), substantial background binding in the HLA64-RC assay requires additional dilution, which can reduce sensitivity. Because the assay is cell-based, surfactants cannot be used during washing to minimize nonspecific binding, and incorporating an IgG purification step may help address this limitation.

We integrated the HLA64-RC assay with a newly optimized single B-cell culture method [43] to identify allogeneic HLA-specific B cells from sensitized patients. In a pilot experiment (Fig. 6), thirteen HLA-specific BCRs were recovered from the peripheral blood derived from a single sensitized transplant candidate. Phenotypic information was obtained by index sorting, and BCR sequences were determined from clonal cultures by RT-PCR, enabling a linked analysis of phenotype, function, and BCR genetics for each HLA-specific B cell. These results demonstrate that this integrated BCR discovery platform provides an effective and reliable approach for identifying and characterizing HLA-specific B cells, even with HLA probes of limited specificity. Ongoing efforts in our group aim to develop a new format of HLA probes with increased specificity, which may enhance efficiency and maximize the utility of this platform for systematic profiling of allogeneic B-cell responses.

Interesting phenotypic, functional, and BCR genetic characteristics were observed in this small set of thirteen HLA-specific B cells (Fig. 6). All six HLA-specific B cells identified by Panel 2 labeling displayed an IgG^+^ CD24^low^ phenotype. Additional phenotyping markers such as CD20, CD21, CD38, CD11c, CXCR5, FcRL4, and FcRL5 will be required to fully define these cells [64]. Tentatively, within these six IgG^+^ CD24^low^ B cells, two were CD27^+^, a phenotype consistent with activated MBCs, which are known to be elevated in DSA-positive transplant recipients [65]. The remaining four were CD27^low/–^, a phenotype suggestive of atypical, tissue-like, or exhausted MBCs that arise under chronic or repeated antigen exposure [66, 67]. These phenotypic features are compatible with the clinical context, as this patient had experienced AMR and the rejected kidney graft likely served as a persistent source of allo-antigens for two years prior to blood sampling.

The production of rAbs from the identified HLA-specific B cells enabled comprehensive characterization of their binding specificities using both the HLA64-RC and SAB assays. Based on the SAB results, thirteen rAbs exhibited nine distinct binding patterns, indicating substantial diversity in HLA epitope recognition within this small set of Abs. This diversity is consistent with findings from Lund’s group [35], who reported 22 binding patterns among 49 A*01:01-specific rAbs, even when applying a relatively stringent cutoff MFI of 40,000. It is possible that these epitopes are also confined to the same region along the α-helices of the peptide-binding groove, as shown in their study [35], given the superior topographic accessibility and enrichment of polymorphic amino acid residues in this region. However, convergence in paratope footprint does not necessarily indicate convergence in immunogenicity. By definition, distinct binding patterns reflect distinct residue dependencies, as they imply recognition of different sets of polymorphic residues, even when the overall paratope footprints appear similar or partially overlapping. Thus, determining the relative contribution of individual epitopic residues for each BCR or Ab is essential before immunogenicity can be meaningfully defined, particularly for different donor–recipient pairs with different mismatching residues within the shared topographic region.

The combined analysis of rAb binding specificities and BCR genetics yielded an interesting observation. For the two identified B-cell lineages, clonal members exhibited different binding patterns despite sharing the same clonal origin. It is unlikely that members of the same clone recognize entirely unrelated eplets; rather, they most likely target highly overlapping epitopes, with altered dependence on specific epitopic residues as a consequence of somatic hypermutation (SHM) modifying paratope residues. This finding highlights an important limitation of current approaches that infer HLA eplets solely from allele binding patterns. Similar phenomena have been observed in infectious disease settings, where clonally related Abs against influenza hemagglutinin [68], HIV envelope [69], or SARS-CoV-2 receptor-binding domain [70] recognize overlapping epitopes yet display distinct strain- or variant-specific binding patterns due to SHM-altered paratope contacts. While viral antigen variants can influence the selection and shaping of SHM [69], it remains unclear whether self-HLAs or undocumented allo-sensitization events drive the development of alloreactive “breadth”, or whether this breadth instead reflects stochastic mutations employed by the immune system to diversify the memory repertoire for recognizing potential variants [71]. In either case, such events would be expected to increase calculated panel reactive antibody (cPRA) scores, potentially reducing the likelihood of identifying compatible donors for sensitized transplant candidates. Accordingly, evaluating the relative capacity of individual eplet mismatches to induce alloreactive breadth may provide clinically relevant information to guide risk stratification in donor–recipient pairing.

Another notable observation from the BCR genetics analysis is the recurrent Ig V-gene usage among the identified HLA-reactive B-cell clones, illustrated by five clones sharing a single IGHV gene segment and seven of nine clones using the same IGKV allele. This suggests that germline-encoded binding determinants, analogous to the GRAB motifs previously implicated in recognition of pathogen-derived immunodominant public epitopes [59], may also contribute to allogeneic HLA recognition. This possibility has important implications for evaluating the immunological risk of specific eplet mismatches. Eplets that coincide with germline-recognized core motifs may exhibit broader immunodominance across individuals with diverse HLA backgrounds, thereby potentially representing higher-risk mismatches. Further BCR specificity and genetics studies in larger patient cohorts will be essential to confirm this phenomenon and determine its clinical relevance.

In summary, we have established a cost-efficient, multiplex single-HLA binding assay and demonstrated that its integration with an optimized BCR discovery platform provides an effective approach for identifying allogeneic HLA-specific B cells from sensitized transplant candidates. Our pilot study validated the feasibility of this workflow and generated observations with both mechanistic and clinically relevant implications. Broader application of this approach to systematically profile alloreactive B-cell responses should advance our understanding of the molecular basis of allorecognition, facilitate the identification of immunodominant HLA eplets, and ultimately inform immunological risk stratification to improve outcomes in transplant recipients.

## METHODS

### Human serum and PBMC samples

Previously collected, banked, and de-identified serum samples from allo-sensitized transplant candidates were used for *in vitro* antibody binding assays under Duke University Institutional Review Board approval (IRB Pro00104220). In addition, one previously collected, banked, and de-identified PBMC sample from an allo-sensitized transplant candidate was used for *in vitro* single B-cell sorting and culture experiments under IRB Pro00073872. The individual had a history of sensitizing exposures, including pregnancy, blood transfusion, and a prior kidney transplant from an HLA-A*01–mismatched donor that subsequently failed approximately two years prior to PBMC collection (consistent with serum reactivity to A*01:01).

### Plasmids, DNA fragments, and gene cloning

Lentiviral transfer vector plasmids pLB-EF1a and pLB-EXIP were generated previously [41]. pMD2.G (Addgene plasmid #12259) and psPAX2 (Addgene plasmid #12260) were gifts from Didier Trono. Double-stranded DNA fragments encoding the following genes were synthesized (Gene Universal Inc. or Twist Bioscience) based on their published amino acid sequences, including EBFP2 [47], mTurquoise2 [48], LSSmOrange [49], hmKeima8.5 [50], NowGFP [51], mKelly1 [52], the neomycin-resistance gene (GenBank V00618.1, with R177S mutation), the hygromycin-resistance gene (UniProt P00557), human CD74 (GenBank NM_001025159.2), and 73 unique HLA alleles (IPD-IMGT/HLA Database [72]) listed in Fig. 1a.

Standard molecular cloning procedures were used for plasmid modification and gene cloning. Lentiviral transfer vector plasmids pLB-EXIN and pLB-EXIH were constructed by replacing the puromycin-resistance gene in pLB-EXIP with the neomycin- or hygromycin-resistance gene, respectively. The six FP genes were cloned into pLB-EF1a, and the human CD74 gene was cloned into pLB-EXIH. HLA allele genes from the HLA-A, -B, -DQA1, and -DRB1/3/4/5 loci were cloned into pLB-EXIP, and those from the DQB1 and DRA loci into pLB-EXIN. Endotoxin-free plasmid DNA was prepared using the E.Z.N.A.® Endo-free Plasmid DNA Mini Kit II (Omega Bio-tek) for mammalian cell transfection. All recombinant plasmid sequences were verified by Sanger DNA sequencing (Genewiz).

### Abs and cell labeling reagents

Fluorochrome-conjugated HLA-specific mAbs used in this study included: PE-conjugated clone W6/32 (W6/32-PE, class I pan-specific, BioLegend 311430), REA274-PE (Bw4 pan-specific, Miltenyi 130-123-985), REA143-PE (Bw6 pan-specific, Miltenyi 130-128-123), REA303-PE (DQ pan-specific, Miltenyi 130-123-765), HLADQ1-PE (DQ pan-specific, BioLegend 318106), SK10-APC (DQ pan-specific, ThermoFisher 17-9881-42), Tu39-PE (class II pan-specific, BioLegend 361716), L243-PE (DR pan-specific, BioLegend 307606), and NFLD.D1-PE (DR4-specific, BD 571189). Fluorochrome-unconjugated HLA-specific mAbs included: W6/32-hIgG1K (W6/32 in human IgG1 format, homemade [42]), BB7.2-hIgG1K (BB7.2 in human IgG1 format, A*02-specific, homemade [42]), GAP.A3 (A*03-specific, homemade [42]), BB7.1 (B*07-specific, BioLegend 372402), Tu169 (DQ pan-specific, BioLegend 361502), and REA332-biotin (class II pan-specific, Miltenyi 130-104-869). Isotype control Abs included human IgG1κ (hIgG1K, Southern Biotech 0151K-01) and human IgG1λ (hIgG1L, Southern Biotech 0151L-01). Secondary labeling reagents included PE-conjugated anti-human IgG (anti-hIgG-PE, 2 µg/ml, Southern Biotech 2040-09), PE-conjugated anti-mouse IgG (anti-mIgG-PE, 2 µg/ml, Southern Biotech 1030-09), and PE-conjugated streptavidin (SA-PE, 2 µg/ml, BD 554061).

Abs and labeling reagents used for labeling of B cells enriched from patient-derived PBMCs included: biotinylated HLA-A*01:01, A*02:01 and A*11:01 monomers (HLA Protein Technologies Inc.), IgG-BB515 (clone G18-145, BD 564581), IgD-PE-Cy7(clone IA6-2, BioLegend 348210), CD19-APC-Fire750 (clone HIB19, BioLegend 302258), CD27-BV421 (clone M-T271, BioLegend 356418), IgM-BV510 (clone MHM-88, BioLegend 314522), CD3-BV650 (clone UCHT1, BioLegend 300468), CD14-BV650 (clone M5E2, BioLegend 301836), CD16-BV650 (clone 3G8, BioLegend 302042), CD24-BV711 (clone ML5, BioLegend 311136), SA-PE (BD 554061), and 7-AAD (BD 559925). Two phenotyping Ab panels were tested in this pilot experiment. Panel 1 included Abs against Dump markers (CD3/CD14/CD16), CD19, and IgD, along with SA-PE and 7-AAD. Panel 2 expanded on this by adding Abs against IgG, IgM, CD27, and CD24.

### Culture of mammalian cell lines

The K530 cell line was generated previously [41] from the K562 cell line (ATCC CCL-243) by knockout of the human *CD32A* gene using standard CRISPR-Cas9 technology. K530 and derivative cell lines were maintained in RPMI-1640 medium (Gibco) supplemented with 10% fetal bovine serum (FBS; Sigma), 10 mM HEPES buffer (Gibco), 55 μM 2-mercaptoethanol (2-ME; Gibco), and 1× Pen-Strep (100 units/ml penicillin and 100 μg/ml streptomycin; Gibco). HEK293T cells (ATCC CRL-11268) were maintained in DMEM (high glucose; Gibco) supplemented with 10% FBS (Sigma), 10 mM HEPES buffer, and 55 μM 2-ME. Expi293F™ cells (Gibco) were maintained in Expi293 Expression Medium (Gibco) supplemented with 1× Pen-Strep.

MEC-147 is an optimized feeder cell line for supporting single B-cell cultures from both humans and rhesus macaques. Briefly, the parental MS40L-low cell line [58] was engineered to autonomously express cytokines required for B-cell proliferation and differentiation, including human IL-2, IL-4, IL-21, and BAFF. Similar cloning efficiencies and Ab production were achieved compared with the parental MS40L-low subline supplemented with exogenous cytokines [43]. MEC-147 cells were maintained in IMDM (Gibco) supplemented with 10% FBS (HyClone) and 55 μM 2-ME.

### Reporter cell line generation and stock vial preparation

Reporter cell lines were generated using procedures identical to those previously reported [41]. Briefly, lentiviral transfer vector plasmids were co-transfected into HEK293T cells with the packaging plasmids pMD2.G and psPAX2 using Lipofectamine 3000 Transfection Reagent (Invitrogen). K530 and derivative cell lines were transduced with filtered lentiviral supernatants by spinoculation at 1,000 × g for 45 min at 32 °C. Following selection with the corresponding antibiotics, monoclonal cell lines were established by single-cell sorting.

Stock vials were prepared by pooling 10–20 individual reporter cell lines to generate each sub-panel. Four sub-panels were assembled to constitute the 64-plex reporter cell panel. The ratios of individual cell lines in these sub-panels were prospectively adjusted to achieve approximately equal proportions after four days of expansion of the final 64-plex panel.

### High-throughput HLA64-RC assay

Pooled and expanded 64-plex reporter cells were aliquoted into 96-well V-bottom plates at 0.1–0.15 × 10^6^ cells per well. After centrifugation and removal of supernatants, primary Ab-containing samples, including sera, single B-cell culture supernatants, or rAbs, were added and incubated for 30 min at 4 °C in the dark. Following two washes with labeling buffer (2% FBS in PBS), secondary Abs were added and incubated for an additional 30 min at 4 °C in the dark. After two further washes, cells were resuspended in 40 µl/well of labeling buffer and analyzed on flow cytometers equipped with high-throughput samplers.

### Design and development of the *HLA64* R package

The *HLA64* R package was designed not as a generalizable and comprehensive flow cytometry data analysis tool, but rather specifically dedicated to the analysis of data derived from the HLA64-RC assay. It focuses on batch processing and visualization, and utilizes a deliberately simplified workflow for efficiency that is robust and well suited within the scope of this assay.

Standard preprocessing steps, including compensation and logicle transformation, were built upon the *flowCore* R package. Data processing, including population subsetting and MFI calculation, was implemented using the *data.table* framework for computational efficiency. Data visualization was performed using the *ggplot2* package. Gating of the live-cell population and determination of cutoff values to separate negative and positive populations for FP channels were performed using custom R functions employing simplified approaches dedicated to the HLA64-RC assay. The robustness of these simplified approaches was safeguarded through user-guided parameter settings and visual verification using exported plots. The full implementation is provided in the deposited R package (see Code Availability).

For live gating, debris and dead cells were first excluded using user-defined forward and side scatter (FSC/SSC) cutoffs, after which the remaining events formed a single dominant population in FSC/SSC space for the HLA64-RC assay. Under these assay-specific conditions, a two-dimensional kernel density estimation (KDE) was applied to define a density-based gating region capturing the majority of events, based on a user-defined percentage (default 85%). Parameters including FSC/SSC cutoffs, KDE grid resolution, and the target coverage percentage are adjustable. Gating plots for individual samples were exported for visual verification.

For FP cutoff determination, the FP-positive and negative populations consistently formed the two dominant populations across all six FP channels in the HLA64-RC assay. Under these conditions, a two-component Gaussian mixture model was fit to the signal distribution after trimming of extreme values. The intersection of the two modeled components was used as the primary cutoff. When substantial overlap was detected at the intersection, a local minimum (“valley”) in the one-dimensional KDE was used as an alternative cutoff. Adjustable parameters include the proportion of signal trimming and the overlap threshold used to trigger KDE-based valley detection. Cutoff plots for individual FP channels were again exported for visual verification.

### Analysis of HLA64-RC assay results

Flow cytometry data from HLA64-RC assays were analyzed using FlowJo™ software or the HLA64 R package. When FlowJo™ was used, live-cell gating and fluorescent protein (FP) cutoff thresholds were defined manually using a control sample and applied across batches. When the HLA64 R package was used, live-cell gating and FP cutoff determination were performed using automated procedures based on a limited set of user-defined parameters, as detailed above. In both cases, the 64 cell populations were subsequently demultiplexed based on the FP cutoffs.

For both FlowJo™- and HLA64-based analyses, positive binding was defined using combined intra-sample and inter-sample thresholds based on exported Ab-binding MFI values for individual demultiplexed reporter cell populations. Binding for a given reporter cell line was considered positive if the MFI value exceeded both (1) the corresponding negative control reporter cell line within the same well and (2) the MFI value for the same reporter cell line from an external control sample or dataset, by at least 40 MFI units or the control MFI value, whichever was greater. The external control sample or dataset was defined either by a negative labeling control sample or by the median MFI value for each reporter cell line across a batch, based on the assumption that, in a typical screening experiment, fewer than half of samples within a batch exhibit positive binding to a given HLA allele. Only events meeting both criteria were classified as positive. This definition was visually verified using overlaid histograms, in which positive populations displayed a clear shift relative to negative controls. The full implementation is provided in the deposited R code (see Code Availability).

### HLA SAB assay and related data analysis

Clinical HLA SAB antibody testing was performed according to manufacturer’s instructions using LABScreen Single Antigen kits (One Lambda, ThermoFisher). Sera were EDTA pretreated, and rAbs were diluted in assay buffer to the indicated concentrations. Assays were acquired on a Luminex 3D instrument (Luminex).

Positive binding thresholds for SAB data were defined empirically based on a small set of patient serum samples (Fig. 4; n = 10). Each serum sample was tested independently, and no shared negative control samples were available. Consequently, thresholds combining normalized bead signals, allele-specific background estimates, and absolute MFI cutoffs were chosen to capture clear positive signals while minimizing false positives. For each sample, the MFI of negative control beads was used to calculate the normalized background (NBG) for each HLA allele (NBG = MFI / negative control bead MFI). Two complementary criteria were applied to identify positive binding: (1) a fixed ratio threshold, where NBG > 6, and (2) an allele-specific threshold. The allele-specific threshold for each HLA allele was derived from the lower half of NBG values across all samples and calculated as the median plus six times the median absolute deviation (MAD). Signals were further required to exceed an absolute MFI of 50 units. Binding to a given HLA allele bead was considered positive if it met the absolute MFI criterion and satisfied either the fixed ratio or allele-specific threshold. For consistency, the same definition was applied to SAB testing with rAb samples (Fig. 6g; n = 13). These thresholds provide a reproducible, data-driven approach for this study, though adjustment may be necessary for other datasets or larger cohorts. We note that, unlike clinical SAB interpretation, we did not apply additional eplet- or pattern-based filtering; near-threshold signals are therefore reported transparently and intended only for technical comparison with the HLA64-RC assay. The full implementation is provided in the deposited R code (see Code Availability).

### Single-B-cell sorting and culture

Frozen PBMCs derived from sensitized transplant candidates were thawed, and B cells were enriched using the Dynabeads™ Untouched Human B Cell Kit (Invitrogen 11351D). HLA monomer probes were selected based on serum reactivity determined by clinical SAB assays. A cocktail of selected biotinylated HLA monomers (final concentration, 1 µg/mL each) was added to enriched B cells and incubated at 4°C for 30 min in the dark. Following one wash with B-cell culture medium (BCM; RPMI-1640 supplemented with 10% FBS (HyClone), 10 mM HEPES, 1 mM sodium pyruvate (Gibco), 1× MEM NEAA (Gibco), 55 µM 2-ME, and 1× Pen-Strep), cells were labeled with phenotyping Ab panels at 4°C for 30 min in the dark. After an additional wash with BCM, cells were resuspended in BCM and kept on ice until single-B-cell sorting.

Single-cell sorting was performed on a BD Aria II at the Duke Human Vaccine Institute Research Flow Cytometry Facility. 7-AAD^−^ Dump^−^ CD19^+^ IgD^−^ SA^+^ B cells were gated and index-sorted into 96-well flat-bottom culture plates containing MEC-147 feeder cell monolayers, which had been pre-seeded 24 h prior to sorting at 3,000 cells per well in BCM. Culture medium was replaced with fresh BCM every 3–4 days. Culture supernatants were harvested on day 25, and the drained culture plates containing expanded B cells and PCs were stored at -80°C for subsequent RNA extraction.

### Human Ig ELISA assay

Single-B-cell culture supernatants were screened for the presence of human Ig using standard sandwich ELISA assays. Briefly, 384-well high-binding ELISA plates (Corning, 3700) were coated with anti-human Igκ (Southern Biotech, 2061-01) and anti-human Igλ (Southern Biotech, 2071-01) Abs at 1 µg/mL each in carbonate buffer. After three washes and blocking, culture supernatants were added at a 1:100 dilution. Following three additional washes, a cocktail of HRP-conjugated anti-human Igκ (Southern Biotech, 2061-05) and anti-human Igλ (Southern Biotech, 2071-05) Abs was added. After three further washes, TMB substrate (BioLegend, 421101) was added and incubated at room temperature in the dark for 5 min. The reaction was stopped by addition of stop solution (1 M H_2_SO_4_), and optical density (OD) values were measured at 450 nm with background subtraction at 650 nm. Supernatants from culture wells without sorted B cells served as negative controls. Culture supernatants with OD values exceeding the mean plus six times the standard deviation (mean + 6 × SD) of negative control OD values were considered Ig-positive.

### BCR V(D)J cloning and sequencing

RNA was extracted from the corresponding original culture plate wells whose supernatants demonstrated HLA-binding activity. BCR V(D)J rearrangements were amplified as previously described [58] and cloned in-frame into pcDNA3.1 (Invitrogen)–derived expression vectors containing human IgG1, Igκ, or Igλ constant regions. Endotoxin-free plasmids were prepared using the E.Z.N.A.® Endo-free Plasmid DNA Mini Kit II (Omega Bio-tek, Cat# D6950). V(D)J sequences were determined by Sanger DNA sequencing at the Duke DNA Sequencing Facility.

### rAb production and purification

Expi293F™ cells were co-transfected with paired heavy- and light-chain expression plasmids using the ExpiFectamine™ 293 Transfection Kit (ThermoFisher). Transfected cells were cultured in Expi293™ Expression Medium with shaking at 150 rpm for 72–96 h. rAbs expressed as human IgG1 were purified from culture supernatants using Pierce™ Protein G UltraLink™ Resin (Thermo Scientific, Cat# 53126) and eluted with IgG Elution Buffer (ThermoFisher, Cat# 21004). Eluted rAbs were buffer-exchanged into PBS using Amicon® Ultra-4 centrifugal filter units (Millipore, Cat# UFC805096), sterilized by filtration through Millex-GV syringe filters (Millipore, Cat# SLGV013SL), and quantified using a NanoDrop 2000 spectrophotometer (ThermoFisher).

### BCR V(D)J clonal analysis

V(D)J gene assignment for both heavy- and light-chain sequences of cloned HLA-specific BCRs was performed using *IgBLAST* [73], based on human IGV, IGD, and IGJ germline databases downloaded from IMGT®, the international ImMunoGeneTics information system® [74]. Clonal relationships were inferred from *IgBLAST*-assigned heavy-chain sequences using the *scoper* program [75] with a hierarchical clustering-based model [76] and an arbitrary distance threshold of 0.2. Germline heavy-chain sequences were inferred using the *Change-O* toolkit [77] via the *CreateGermlines.py* program, and phylogenetic trees were reconstructed using *BuildTrees.py* and *IgPhyML* [78]. Phylogenetic trees of identified HLA-specific B-cell lineages were visualized using the R package *ggtree* [79]. The full implementation is provided in the deposited R and Python code (see Code Availability).

### Software and computer programs used for data processing, analysis and visualization

Flow cytometry data were analyzed using either FlowJo™ or the *HLA64* R package (v0.1.0) developed in this study (see Code Availability). BCR V(D)J sequencing data were analyzed using *IgBlast* (v1.22.0; https://ftp.ncbi.nih.gov/blast/executables/igblast/release/1.22.0/), the R package *scoper* (v1.3.0), and the Python packages *Change-O* (v1.3.4) and *IgPhyML* (v2.1.0). Data from other assays were preprocessed in Microsoft Excel (v1808). All data (except flow cytometry data) were processed, analyzed, and visualized using R (v4.5.2) and RStudio (v2025.09.2). Final figures were compiled with Adobe Illustrator (v29.7.1).

The *HLA64* R package was developed using the following R packages: *callr* (v3.7.6), *data.table* (v1.17.8), *dplyr* (v1.1.4), *egg* (v0.4.5), *filelock* (v1.0.3), *flowCore* (v2.20.0), *furrr* (v0.3.1), *future* (v1.68.0), *ggplot2* (v4.0.1), *ggridges* (v0.5.7), *graphics* (v4.5.2), *grDevices* (v4.5.2), *grid* (v4.5.2), *gridExtra* (v2.3), *gtools* (v3.9.5), *magrittr* (v2.0.4), *MASS* (v7.3.65), *mixtools* (v2.0.0.1), *openxlsx* (v4.2.8.1), *parallel* (v4.5.2), *patchwork* (v1.3.2), *purrr* (v1.2.0), *scales* (v1.4.0), *shiny* (v1.12.1), *shinyFiles* (v0.9.3), *sp* (v2.2.0), *stats* (v4.5.2), *stringr* (v1.6.0), *tidyr* (v1.3.1), *tools* (v4.5.2), and *utils* (v4.5.2). Additional R packages used for data analysis and visualization included *ape* (v5.8.1), *Biostrings* (v2.76.0), *cowplot* (v1.2.0), *DescTools* (v0.99.60), *ggtree* (v3.16.3), *ggVennDiagram* (v1.5.4), *lmodel2* (v1.7.4), and *tidyverse* (v2.0.0).

### Statistical analysis

Agreement between HLA64-RC and SAB assay measurements was assessed using symmetric regression and concordance metrics. Standardized major axis (SMA) regression was performed on log_10_-transformed MFI values using the *lmodel2* R package to account for measurement error in both variables. Lin’s concordance correlation coefficient (CCC) was calculated, and 95% confidence intervals for SMA slope, intercept, and CCC were estimated by non-parametric bootstrap resampling (1,000 iterations). Spearman’s rank correlation coefficient was calculated as a complementary, distribution-independent measure of association. For comparison of data processing pipelines, paired MFI values generated by HLA64 and FlowJo™ were compared using ordinary least squares linear regression, with Pearson’s and Spearman’s correlation coefficients reported for descriptive comparison. All analyses were performed in R (see Code Availability).

## DATA AVAILABILITY

V(D)J sequences derived from the thirteen identified HLA-specific BCRs have been deposited in NCBI GenBank under accession numbers PX727065–PX727090. Source data underlying the figures, including data presented post-processing or indirectly, have been deposited in Zenodo under DOI 10.5281/zenodo.18291870. All other data supporting the findings of this study are available from the corresponding author upon reasonable request.

## CODE AVAILABILITY

The *HLA64*R package is publicly available on GitHub (https://github.com/shengli-song/HLA64) under the Artistic License 2.0. The full package documentation and vignettes are available via the GitHub repository. Custom R and Python scripts used for data processing, analysis, and visualization have been deposited in Zenodo together with the corresponding source data under the same DOI as listed in the Data Availability statement.

## ACKNOWLEDGEMENTS

We thank Xiaoyan Nie, Xiaoe Liang, and Dongmei Liao for technical assistance. This study was supported by grants AI163065 (NIAID), ITN089ST (Benaroya Research Institute), and C2833376 (Haller Foundation) to S.J.K.

## AUTHOR CONTRIBUTIONS

S.J.K. obtained funding and supervised the study. S.S. conceived and designed the study. S.S., E.F.K., and A.M.D. performed the experiments. G.K. provided crucial materials. A.M.J. identified patient samples and contributed to data interpretation. S.S. developed the computational workflow, analyzed and visualized the data, and drafted the manuscript. J.K., C.C., A.M.J., G.K., and S.J.K. analyzed data and contributed consultation and scientific discussion. All authors reviewed and revised the manuscript.

---

**Supplementary Fig. 1.**
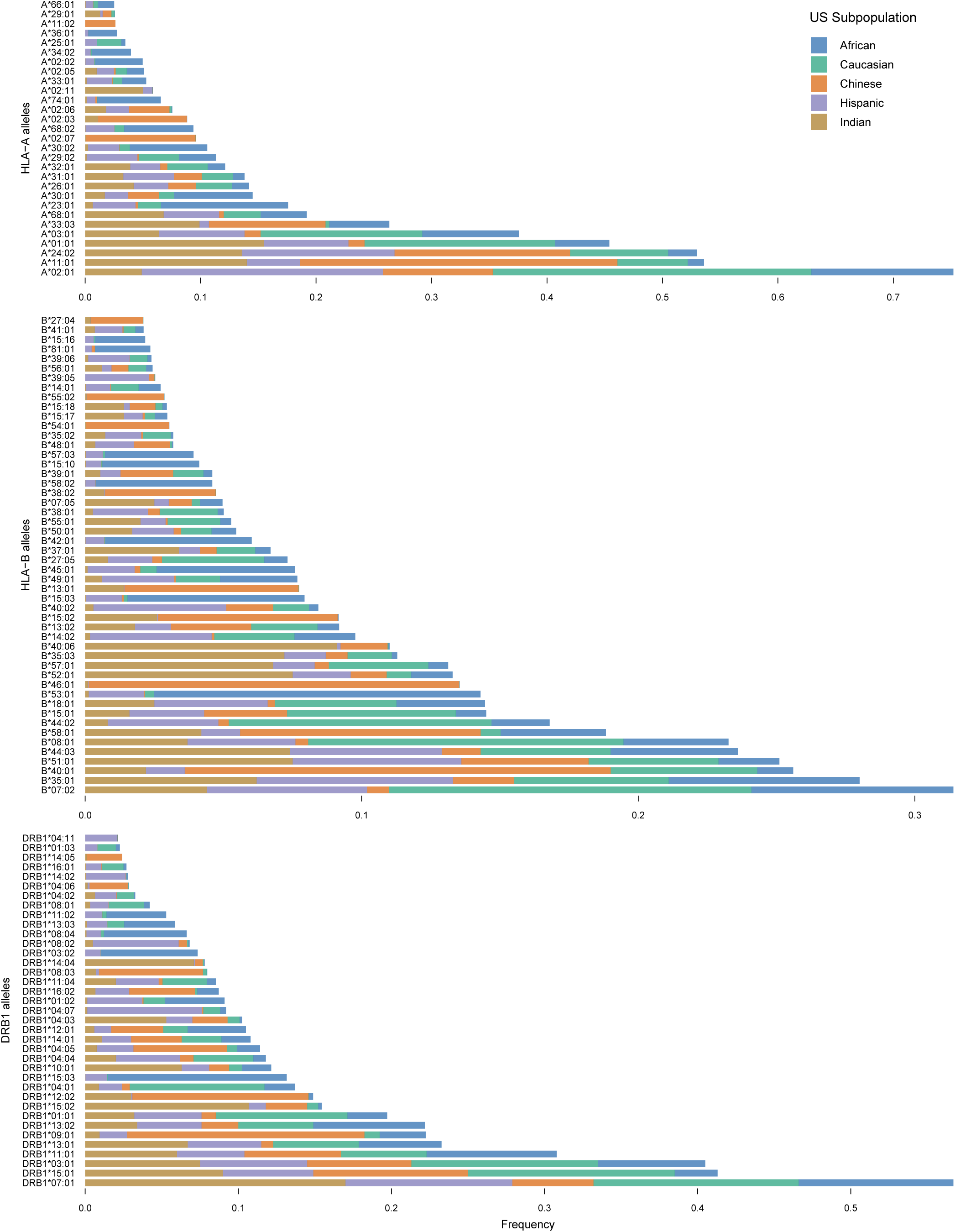
Top frequent HLA-A, -B, and -DRB1 alleles across five genetically diverse U.S. subpopulations. HLA allele frequency data were retrieved from the Allele Frequency Net Database. Records from the National Marrow Donor Program (NMDP) were used for five genetically diverse U.S. subpopulations with large sample sizes (Supplementary Date 1), including African American pop 2 (n = 416,581), European Caucasian (n = 1,242,890), Chinese (n = 99,672), Hispanic South or Central American (n = 146,714), and South Asian Indian (n = 185,391). For each HLA locus (HLA-A, -B, and -DRB1), alleles were ranked by their cumulative frequencies across the five subpopulations. Only alleles with cumulative frequency greater than 0.02 are shown.

**Supplementary Fig. 2.**
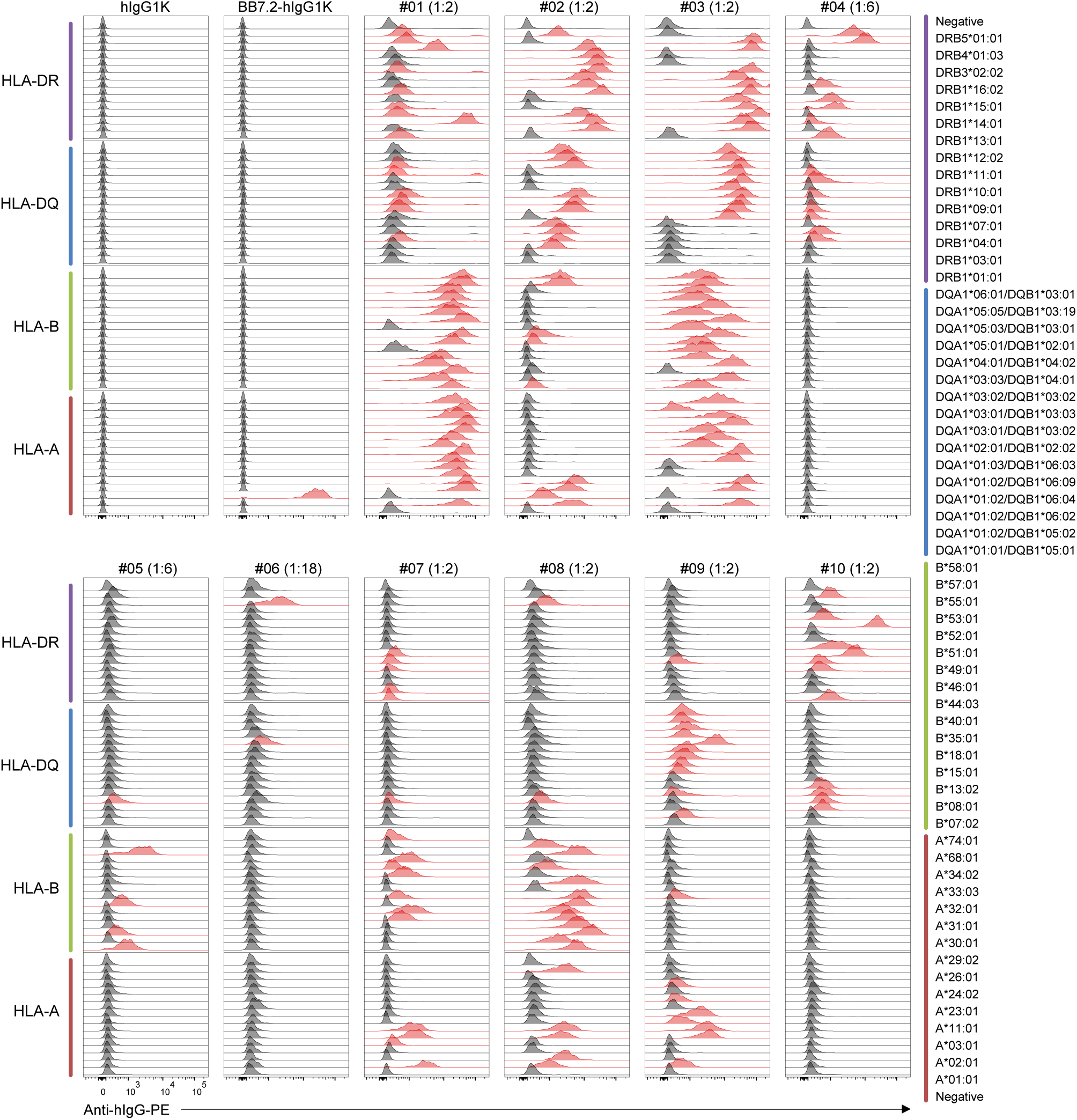
HLA-specific serum IgG binding activities detected using the HLA64-RC assay at selected dilutions. Serum samples from ten sensitized transplant candidates (#01–#10) were analyzed in HLA64-RC assays as detailed in Supplementary Document 1. For each serum, the dilution (indicated in parentheses) yielding the highest signal-to-background ratio was selected for MFI calculation (Supplementary Data 3) and visualization. Populations with MFI values above the binding threshold (see Methods) are highlighted in red. Histograms are grouped by HLA-A (dark red), HLA-B (green), HLA-DQ (blue), and HLA-DR (purple) alleles, displayed upward in the order indicated on the far-right. Control primary antibodies (1 µg/ml) included human IgG1κ isotype control (hIgG1K) and BB7.2 in human IgG1κ format (BB7.2-hIgG1K).

**Supplementary Fig. 3.**
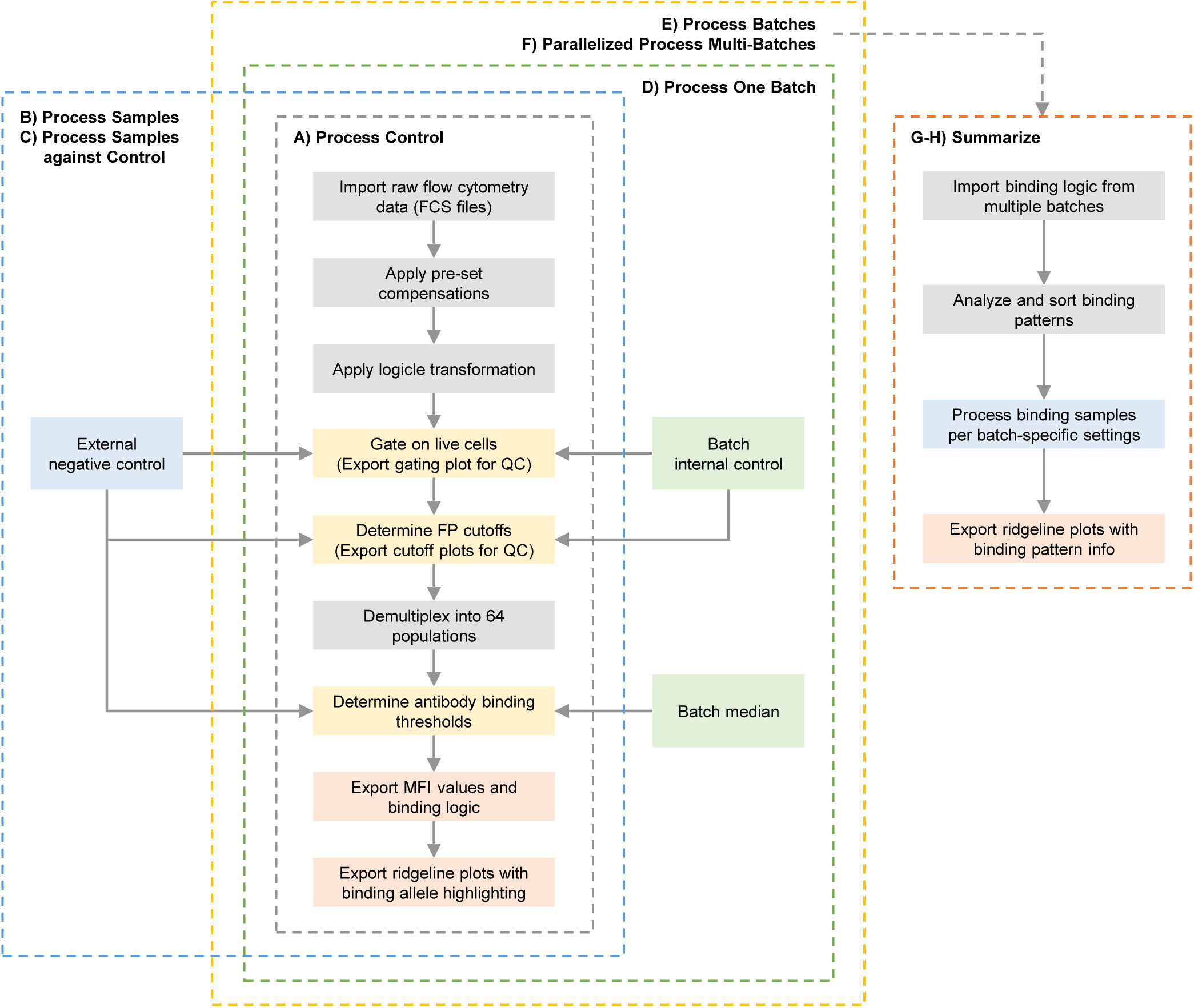
Overall workflow of the HLA64 R package. Raw flow cytometry data are imported from individual FCS files, and pre-set compensation is applied. Logicle transformation is then applied to the six FP channels. Live-cell gating is automatically established based on analysis of an external negative control or a batch-internal control sample and subsequently applied to all samples within the batch. Cutoff values are automatically determined for each FP channel to separate negative and positive populations, enabling demultiplexing of 64 distinct cell populations. These cutoffs are derived from the control sample and applied to all other samples within the same batch. Live-cell gating and FP cutoff plots are exported for visual verification. Ab-binding MFI values are calculated for each demultiplexed population, and binding thresholds are determined by referencing either the MFI values of the external negative control sample or the median MFI within the batch (see Methods). Finally, MFI values and binding logic for individual populations are exported together with ridgeline plots highlighting the binding alleles. The package provides multiple running modes, which can be conveniently accessed via the graphical user interface (GUI) by invoking *launch_GUI()*. These modes include: (A) processing a control sample for visual verification; (B) processing multiple samples based on predefined control settings; (C) processing samples against a control; (D) processing a single batch against an internal control; and (E) processing multiple batches sequentially or (F) in parallel for improved efficiency. In HTS applications, binding patterns from multiple batches can be summarized and analyzed collectively (modes G and H). Binding patterns are sorted based on the exported binding logic, and the corresponding samples are processed using batch-specific settings. Aggregated ridgeline plots and summarized binding pattern data are then exported for downstream analysis.

**Supplementary Fig. 4.**
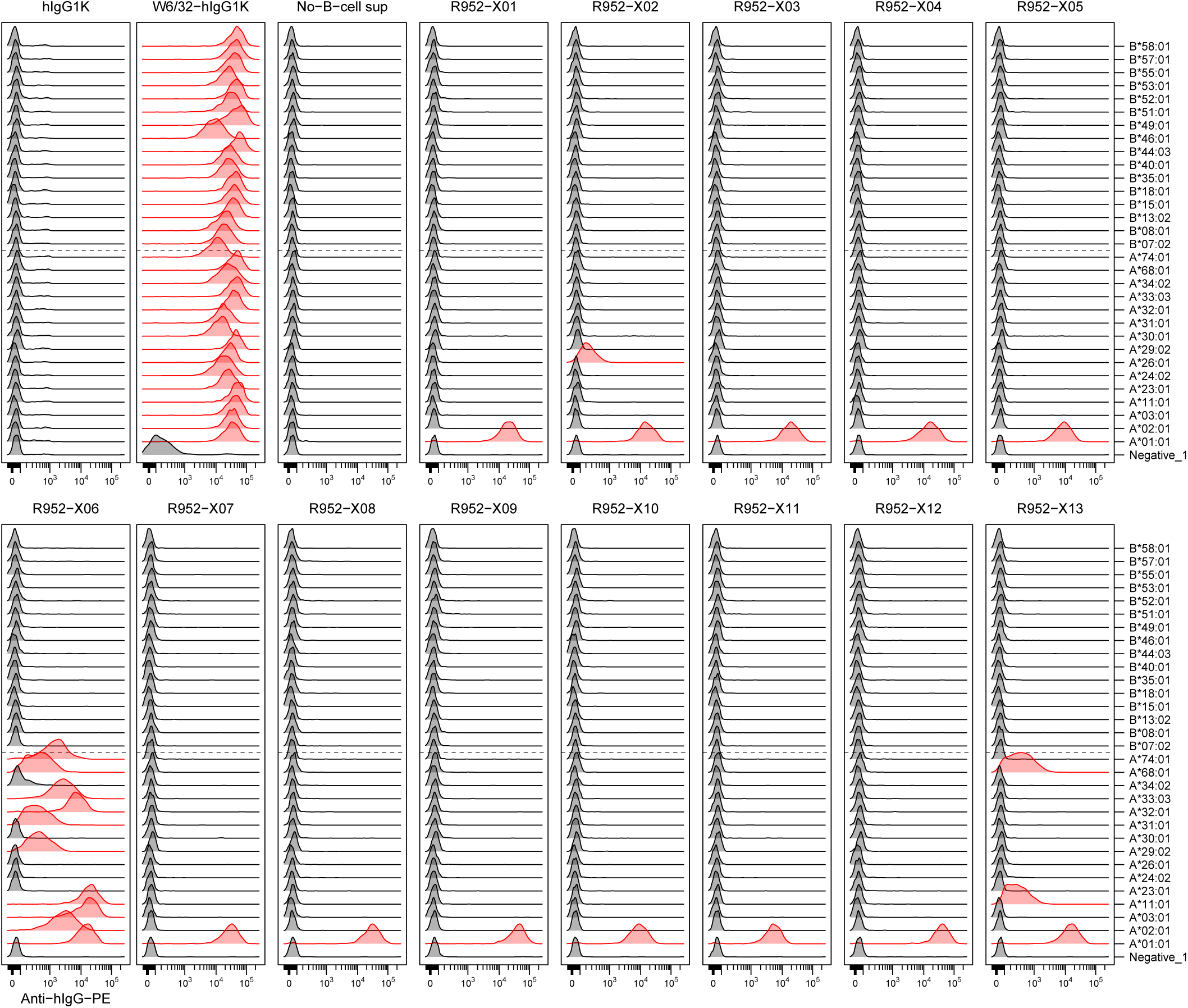
HLA-specific single B-cell cultures identified using the HLA64-RC assay. HLA monomer-binding single B-cell culture supernatants were screened using the HLA64-RC assay. R952-X06 and R952-X08 to -X12 cultures were derived from phenotyping antibody Panel 1 mediated sorting, and the rest from Panel 2 sorting. Secondary labeling was performed using PE-conjugated anti-human IgG (2 μg/ml). Flow cytometry data were analyzed and visualized using the *HLA64* R package, and MFI values were exported (Supplementary Data 5). Demultiplexed HLA alleles corresponding to individual reporter cell populations are indicated on the right. Data are shown for HLA-specific cultures, and only HLA-A and -B loci are plotted. Human IgG1κ (hIgG1K), W6/32 in human IgG1κ format (W6/32-hIgG1K), and a no-B-cell culture supernatant were included as labeling controls. Populations exceeding the binding threshold (see Methods) relative to the no-Bcell control were automatically highlighted in red.

**Supplementary Document 1. HLA-specific serum IgG binding activities detected using the HLA64-RC assay.**

The HLA64-RC panel was labeled with control antibodies and serum samples from ten sensitized transplant candidates (#01–#10), each tested in 1/3 serial dilutions from 1:2 to 1:486 (dilutions indicated in parentheses). Secondary labeling was performed using PE-conjugated anti-human IgG (2 µg/ml). Flow cytometry data were analyzed as described in Fig. 3, and binding activities were plotted accordingly. Histograms are grouped by HLA-A (dark red), HLA-B (green), HLA-DQ (blue), and HLA-DR (purple) alleles, displayed upward in the order indicated on the far-right. Control primary antibodies, including human IgG1κ isotype control (hIgG1K) and BB7.2 in human IgG1κ format (BB7.2-hIgG1K), were tested in 1/3 serial dilutions from 1000 to 4.12 ng/ml (concentrations indicated in parentheses).

**Supplementary Document 2. Binding activities of serial dilutions of HLA-specific rAbs detected using the HLA64-RC assay.**

The HLA64-RC panel was labeled with serial dilutions of HLA-specific rAbs at 5000, 500, 50, 5, and 0.5 ng/ml (concentrations indicated in parentheses), followed by secondary labeling with PE-conjugated anti-human IgG (2 µg/ml). Flow cytometry data were analyzed and visualized using the *HLA64* R package, and MFI values were exported (Supplementary Data 5). Labeling buffer only, human IgG1κ (hIgG1K, 5 µg/ml), human IgG1λ (hIgG1L, 5 µg/ml), and serial dilutions of W6/32 in human IgG1κ format (W6/32-hIgG1K) were included as labeling controls. Populations exceeding the binding threshold (see Methods) relative to the labeling buffer control were automatically highlighted in red.

**Supplementary Data 1.** Allele frequency data used in Fig. 1b, c and Supplementary Fig. 1.

**Supplementary Data 2.** Eplet information used in Fig. 1d.

**Supplementary Data 3.** Serum binding MFI values obtained using the HLA64-RC assay and used in Figs. 4 and 5e.

**Supplementary Data 4.** Serum binding MFI values obtained using the SAB assay and used in Fig. 4.

**Supplementary Data 5.** Antibody binding MFI values obtained using the HLA64-RC assay and used in Fig. 6.

**Supplementary Data 6.** Antibody binding MFI values obtained using the SAB assay and used in Fig. 6.

## REFERENCES

1. Terasaki PI. Humoral theory of transplantation. Am J Transplant 3, 665–673 (2003).

2. Loupy A, Lefaucheur C. Antibody-Mediated Rejection of Solid-Organ Allografts. N Engl J Med 379, 1150–1160 (2018).

3. Sellares J, et al. Understanding the causes of kidney transplant failure: the dominant role of antibody-mediated rejection and nonadherence. Am J Transplant 12, 388–399 (2012).

4. Rodriguez-Ramirez S, Al Jurdi A, Konvalinka A, Riella LV. Antibody-mediated rejection: prevention, monitoring and treatment dilemmas. Curr Opin Organ Transplant 27, 405–414 (2022).

5. Willicombe M, et al. De novo DQ donor-specific antibodies are associated with a significant risk of antibody-mediated rejection and transplant glomerulopathy. Transplantation 94, 172–177 (2012).

6. Mohan S, et al. Donor-specific antibodies adversely affect kidney allograft outcomes. J Am Soc Nephrol 23, 2061–2071 (2012).

7. Bestard O, Couzi L, Crespo M, Kessaris N, Thaunat O. Stratifying the humoral risk of candidates to a solid organ transplantation: a proposal of the ENGAGE working group. Transpl Int 34, 1005–1018 (2021).

8. Kurosaki T, Kometani K, Ise W. Memory B cells. Nat Rev Immunol 15, 149–159 (2015).

9. Mulder A, et al. Identification, isolation, and culture of HLA-A2-specific B lymphocytes using MHC class I tetramers. J Immunol 171, 6599–6603 (2003).

10. Zachary AA, Kopchaliiska D, Montgomery RA, Leffell MS. HLA-specific B cells: I. A method for their detection, quantification, and isolation using HLA tetramers. Transplantation 83, 982–988 (2007).

11. Han M, Rogers JA, Lavingia B, Stastny P. Peripheral blood B cells producing donor-specific HLA antibodies in vitro. Hum Immunol 70, 29–34 (2009).

12. Heidt S, et al. A NOVel ELISPOT assay to quantify HLA-specific B cells in HLA-immunized individuals. Am J Transplant 12, 1469–1478 (2012).

13. Karahan GE, et al. A Memory B Cell Crossmatch Assay for Quantification of Donor-Specific Memory B Cells in the Peripheral Blood of HLA-Immunized Individuals. Am J Transplant 17, 2617–2626 (2017).

14. Luque S, Lucia M, Crespo E, Jarque M, Grinyo JM, Bestard O. A multicolour HLA-specific B-cell FluoroSpot assay to functionally track circulating HLA-specific memory B cells. J Immunol Methods 462, 23–33 (2018).

15. Karahan GE, et al. An Easy and Sensitive Method to Profile the Antibody Specificities of HLA-specific Memory B Cells. Transplantation 103, 716–723 (2019).

16. Snanoudj R, Claas FH, Heidt S, Legendre C, Chatenoud L, Candon S. Restricted specificity of peripheral alloreactive memory B cells in HLA-sensitized patients awaiting a kidney transplant. Kidney Int 87, 1230–1240 (2015).

17. Lucia M, et al. Preformed circulating HLA-specific memory B cells predict high risk of humoral rejection in kidney transplantation. Kidney Int 88, 874–887 (2015).

18. Altulea D, Lammerts RGM, Bungener L, Heeringa P, Berger SP, Sanders JS. Memory B-Cell Assay Uncovers Prior HLA Sensitisation in Transplant Candidates With Prior Exposure to Non-Self HLA. HLA 106, e70393 (2025).

19. Luque S, et al. Value of monitoring circulating donor-reactive memory B cells to characterize antibody-mediated rejection after kidney transplantation. Am J Transplant 19, 368–380 (2019).

20. Duquesnoy RJ. A structurally based approach to determine HLA compatibility at the humoral immune level. Hum Immunol 67, 847–862 (2006).

21. McCaughan JA, et al. Identification of risk epitope mismatches associated with de novo donor-specific HLA antibody development in cardiothoracic transplantation. Am J Transplant 18, 2924–2933 (2018).

22. Hamada S, et al. Predictive value of HLAMatchmaker and PIRCHE-II scores for de novo donor-specific antibody formation after adult and pediatric liver transplantation. Transpl Immunol 61, 101306 (2020).

23. Sapir-Pichhadze R, et al. Epitopes as characterized by antibody-verified eplet mismatches determine risk of kidney transplant loss. Kidney Int 97, 778–785 (2020).

24. Senev A, et al. Eplet Mismatch Load and De Novo Occurrence of Donor-Specific Anti-HLA Antibodies, Rejection, and Graft Failure after Kidney Transplantation: An Observational Cohort Study. J Am Soc Nephrol 31, 2193–2204 (2020).

25. Nguyen HD, et al. Modeling the benefits and costs of integrating an acceptable HLA mismatch allocation model for highly sensitized patients. Transplantation 97, 769–774 (2014).

26. Heidt S, et al. Allocation to highly sensitized patients based on acceptable mismatches results in low rejection rates comparable to nonsensitized patients. Am J Transplant 19, 2926–2933 (2019).

27. Bezstarosti S, et al. A Comprehensive Evaluation of the Antibody-Verified Status of Eplets Listed in the HLA Epitope Registry. Front Immunol 12, 800946 (2021).

28. Bezstarosti S, Kramer CSM, Claas FHJ, de Fijter JW, Reinders MEJ, Heidt S. Implementation of molecular matching in transplantation requires further characterization of both immunogenicity and antigenicity of individual HLA epitopes. Hum Immunol 83, 256–263 (2022).

29. Jucaud V. Allogeneic HLA Humoral Immunogenicity and the Prediction of Donor-Specific HLA Antibody Development. Antibodies (Basel) 13, (2024).

30. Bezstarosti S, Heidt S. The Progress and Challenges of Implementing HLA Molecular Matching in Clinical Practice. Transpl Int 38, 14716 (2025).

31. Kramer CSM, et al. Generation and reactivity analysis of human recombinant monoclonal antibodies directed against epitopes on HLA-DR. Am J Transplant 20, 3341–3353 (2020).

32. Bezstarosti S, et al. HLA-DQ-Specific Recombinant Human Monoclonal Antibodies Allow for In-Depth Analysis of HLA-DQ Epitopes. Front Immunol 12, 761893 (2021).

33. Kramer CSM, et al. Antibody verification of HLA class I and class II eplets by human monoclonal HLA antibodies. HLA 103, e15345 (2024).

34. Duquesnoy RJ, et al. 16th IHIW: a website for antibody-defined HLA epitope Registry. Int J Immunogenet 40, 54–59 (2013).

35. Killian JT, Jr., et al. Topography of the HLA-A protein enforces shared and convergent immunodominant B cell and antibody alloresponses in transplant recipients. Immunity 58, 3040–3060 e3012 (2025).

36. Visentin J, et al. Denatured class I human leukocyte antigen antibodies in sensitized kidney recipients: prevalence, relevance, and impact on organ allocation. Transplantation 98, 738–744 (2014).

37. Lobashevsky AL. Methodological aspects of anti-human leukocyte antigen antibody analysis in solid organ transplantation. World J Transplant 4, 153–167 (2014).

38. Park BG, Park Y, Kim BS, Kim YS, Kim HS. False Positive Class II HLA Antibody Reaction Due to Antibodies Against Denatured HLA Might Differ Between Assays: One Lambda vs. Immucor. Ann Lab Med 40, 424–427 (2020).

39. Karahan GE, de Vaal Y, Bakker K, Roelen D, Claas FHJ, Heidt S. Comparison of different luminex single antigen bead kits for memory B cell-derived HLA antibody detection. HLA 98, 200–206 (2021).

40. Gutierrez-Larranaga M, et al. Detection of antibodies to denatured human leucocyte antigen molecules by single antigen Luminex. HLA 97, 52–59 (2021).

41. Song S, Manook M, Kwun J, Jackson AM, Knechtle SJ, Kelsoe G. A cell-based multiplex immunoassay platform using fluorescent protein-barcoded reporter cell lines. Commun Biol 4, 1338 (2021).

42. Song S, Manook M, Kwun J, Jackson AM, Knechtle SJ, Kelsoe G. Allo-Specific Humoral Responses: New Methods for Screening Donor-Specific Antibody and Characterization of HLA-Specific Memory B Cells. Front Immunol 12, 705140 (2021).

43. Song S, et al. Functional Convergence of Genetically Diverse B-Cell Receptors in Simian-HIV Infected Rhesus Macaques. Preprint at https://www.biorxiv.org/content/10.64898/2026.01.09.698730v1 (2026).

44. Gonzalez-Galarza FF, et al. Allele frequency net database (AFND) 2020 update: gold-standard data classification, open access genotype data and new query tools. Nucleic Acids Res 48, D783–D788 (2020).

45. Creary LE, et al. High-resolution HLA allele and haplotype frequencies in several unrelated populations determined by next generation sequencing: 17th International HLA and Immunogenetics Workshop joint report. Hum Immunol 82, 505–522 (2021).

46. Usureau C, et al. Antibodies against HLA cross-reactivity groups: From single antigen bead assay to immunoinformatics interpretation of epitopes. Mol Immunol 133, 154–162 (2021).

47. Ai HW, Shaner NC, Cheng Z, Tsien RY, Campbell RE. Exploration of new chromophore structures leads to the identification of improved blue fluorescent proteins. Biochemistry 46, 5904–5910 (2007).

48. Goedhart J, et al. Structure-guided evolution of cyan fluorescent proteins towards a quantum yield of 93%. Nat Commun 3, 751 (2012).

49. Shcherbakova DM, Hink MA, Joosen L, Gadella TW, Verkhusha VV. An orange fluorescent protein with a large Stokes shift for single-excitation multicolor FCCS and FRET imaging. J Am Chem Soc 134, 7913–7923 (2012).

50. Guan Y, et al. Live-cell multiphoton fluorescence correlation spectroscopy with an improved large Stokes shift fluorescent protein. Mol Biol Cell 26, 2054–2066 (2015).

51. Sarkisyan KS, et al. Green fluorescent protein with anionic tryptophan-based chromophore and long fluorescence lifetime. Biophys J 109, 380–389 (2015).

52. Wannier TM, et al. Monomerization of far-red fluorescent proteins. Proc Natl Acad Sci U S A 115, E11294–E11301 (2018).

53. Le Bouteiller P, et al. Engagement of CD160 receptor by HLA-C is a triggering mechanism used by circulating natural killer (NK) cells to mediate cytotoxicity. Proc Natl Acad Sci U S A 99, 16963–16968 (2002).

54. Badders JL, Jones JA, Jeresano ME, Schillinger KP, Jackson AM. Variable HLA expression on deceased donor lymphocytes: Not all crossmatches are created equal. Hum Immunol 76, 795–800 (2015).

55. Lachmann N, Todorova K, Schulze H, Schonemann C. Luminex((R)) and its applications for solid organ transplantation, hematopoietic stem cell transplantation, and transfusion. Transfus Med Hemother 40, 182–189 (2013).

56. Parks DR, Roederer M, Moore WA. A new “Logicle” display method avoids deceptive effects of logarithmic scaling for low signals and compensated data. Cytometry A 69, 541–551 (2006).

57. Moore WA, Parks DR. Update for the logicle data scale including operational code implementations. Cytometry A 81, 273–277 (2012).

58. McCarthy KR, et al. Memory B Cells that Cross-React with Group 1 and Group 2 Influenza A Viruses Are Abundant in Adult Human Repertoires. Immunity 48, 174–184 e179 (2018).

59. Shrock EL, et al. Germline-encoded amino acid-binding motifs drive immunodominant public antibody responses. Science 380, eadc9498 (2023).

60. Kosmoliaptsis V, Dafforn TR, Chaudhry AN, Halsall DJ, Bradley JA, Taylor CJ. High-resolution, three-dimensional modeling of human leukocyte antigen class I structure and surface electrostatic potential reveals the molecular basis for alloantibody binding epitopes. Hum Immunol 72, 1049–1059 (2011).

61. Martinino A, et al. A Novel IgG- and IgM-Cleaving Endopeptidase, IceMG, for Antibody-Mediated Rejection. Am J Transplant, (2025).

62. Quon JC, et al. HLA diversity in ethnic populations can affect detection of donor-specific antibodies by single antigen beads. Front Immunol 14, 1287028 (2023).

63. San Segundo D, Comins-Boo A, Lopez-Hoyos M. Anti-Human Leukocyte Antigen Antibody Detection from Terasaki’s Humoral Theory to Delisting Strategies in 2024. Int J Mol Sci 26, (2025).

64. Sanz I, et al. Challenges and Opportunities for Consistent Classification of Human B Cell and Plasma Cell Populations. Front Immunol 10, 2458 (2019).

65. Louis K, et al. T-bet+CD27+CD21-B cells poised for plasma cell differentiation during antibody-mediated rejection of kidney transplants. JCI Insight 6, (2021).

66. Muellenbeck MF, et al. Atypical and classical memory B cells produce Plasmodium falciparum neutralizing antibodies. J Exp Med 210, 389–399 (2013).

67. Holla P, et al. Shared transcriptional profiles of atypical B cells suggest common drivers of expansion and function in malaria, HIV, and autoimmunity. Sci Adv 7, (2021).

68. Sun X, et al. Unique binding pattern for a lineage of human antibodies with broad reactivity against influenza A virus. Nat Commun 13, 2378 (2022).

69. Liao HX, et al. Co-evolution of a broadly neutralizing HIV-1 antibody and founder virus. Nature 496, 469–476 (2013).

70. Mandolesi M, et al. Multi-compartmental diversification of neutralizing antibody lineages dissected in SARS-CoV-2 spike-immunized macaques. Nat Commun 15, 6338 (2024).

71. Korenkov M, et al. Somatic hypermutation introduces bystander mutations that prepare SARS-CoV-2 antibodies for emerging variants. Immunity 56, 2803–2815 e2806 (2023).

72. Barker DJ, et al. The IPD-IMGT/HLA Database. Nucleic Acids Res 51, D1053–D1060 (2023).

73. Ye J, Ma N, Madden TL, Ostell JM. IgBLAST: an immunoglobulin variable domain sequence analysis tool. Nucleic Acids Res 41, W34–40 (2013).

74. Lefranc MP, et al. IMGT(R), the international ImMunoGeneTics information system(R) 25 years on. Nucleic Acids Res 43, D413–422 (2015).

75. Nouri N, Kleinstein SH. Somatic hypermutation analysis for improved identification of B cell clonal families from next-generation sequencing data. PLoS Comput Biol 16, e1007977 (2020).

76. Gupta NT, Adams KD, Briggs AW, Timberlake SC, Vigneault F, Kleinstein SH. Hierarchical Clustering Can Identify B Cell Clones with High Confidence in Ig Repertoire Sequencing Data. J Immunol 198, 2489–2499 (2017).

77. Gupta NT, Vander Heiden JA, Uduman M, Gadala-Maria D, Yaari G, Kleinstein SH. Change-O: a toolkit for analyzing large-scale B cell immunoglobulin repertoire sequencing data. Bioinformatics 31, 3356–3358 (2015).

78. Hoehn KB, Lunter G, Pybus OG. A Phylogenetic Codon Substitution Model for Antibody Lineages. Genetics 206, 417–427 (2017).

79. Yu G, Smith DK, Zhu H, Guan Y, Lam TT. ggtree: an r package for visualization and annotation of phylogenetic trees with their covariates and other associated data. Methods Ecol Evol 8, 28–36 (2017).

