## Supplementary figures and images for "Profiling Allogeneic HLA-specific B-cell Responses Utilizing a 64-plex Single-HLA Reporter Cell Panel"

### Supplementary Document 1

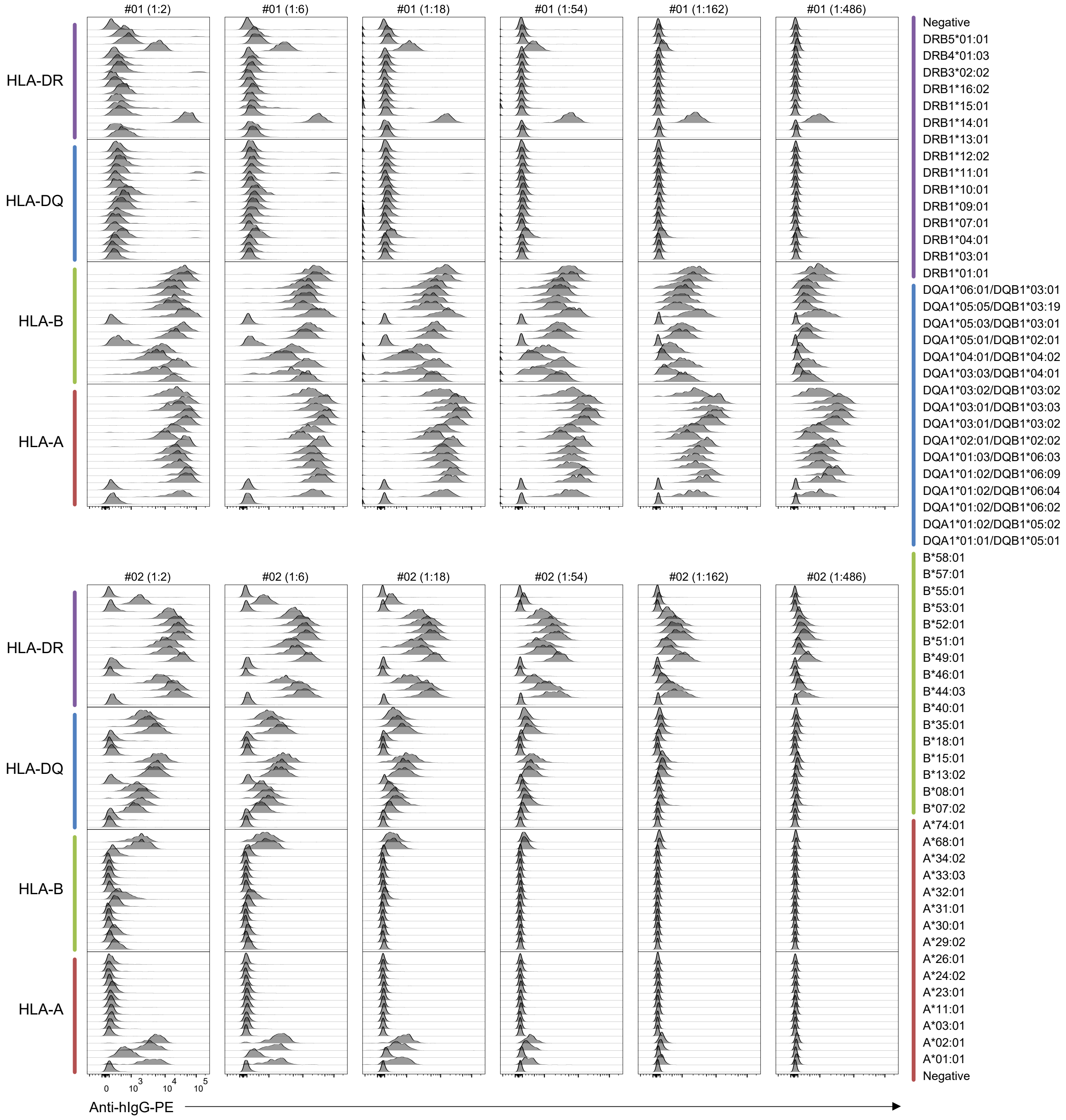

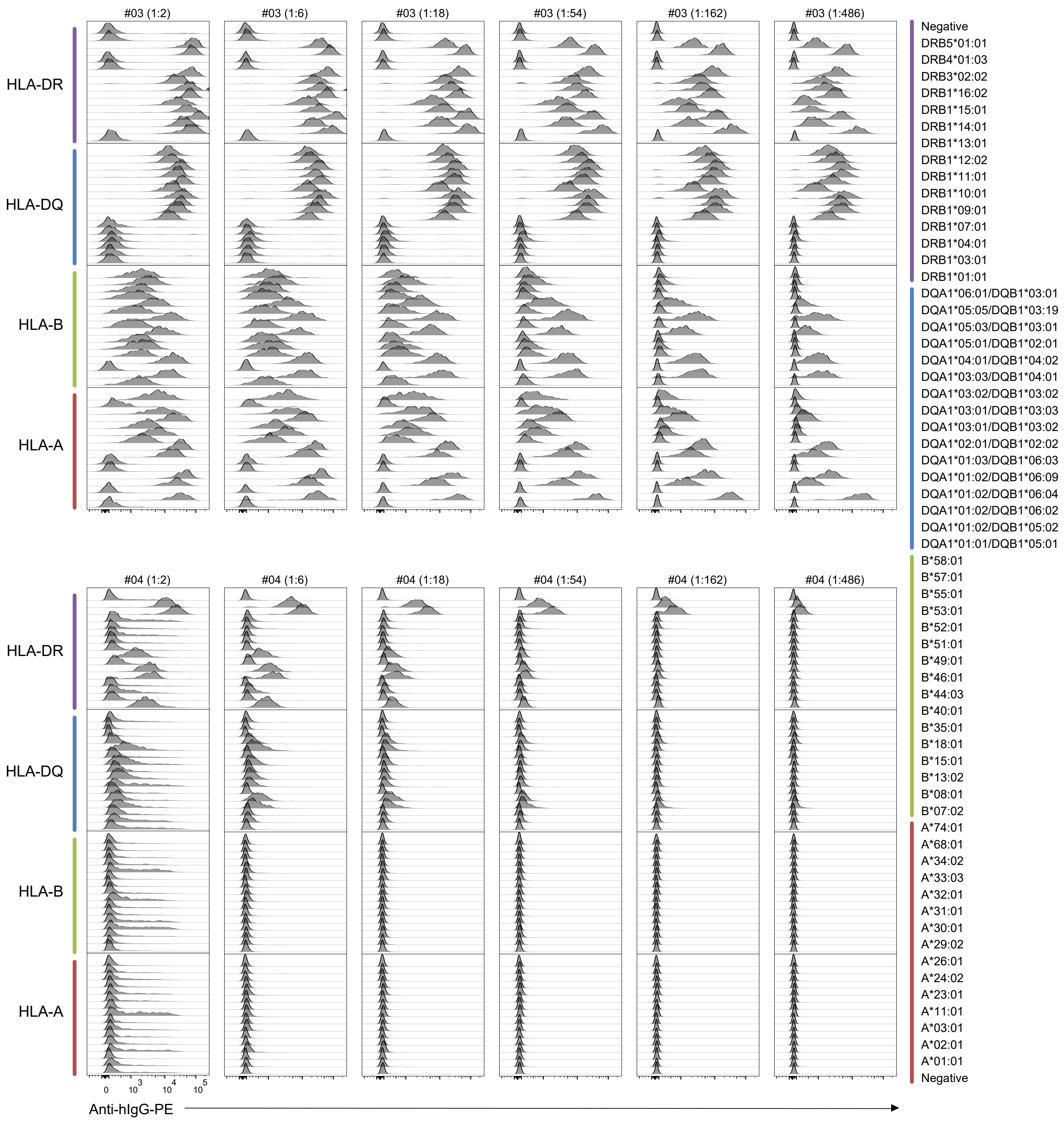

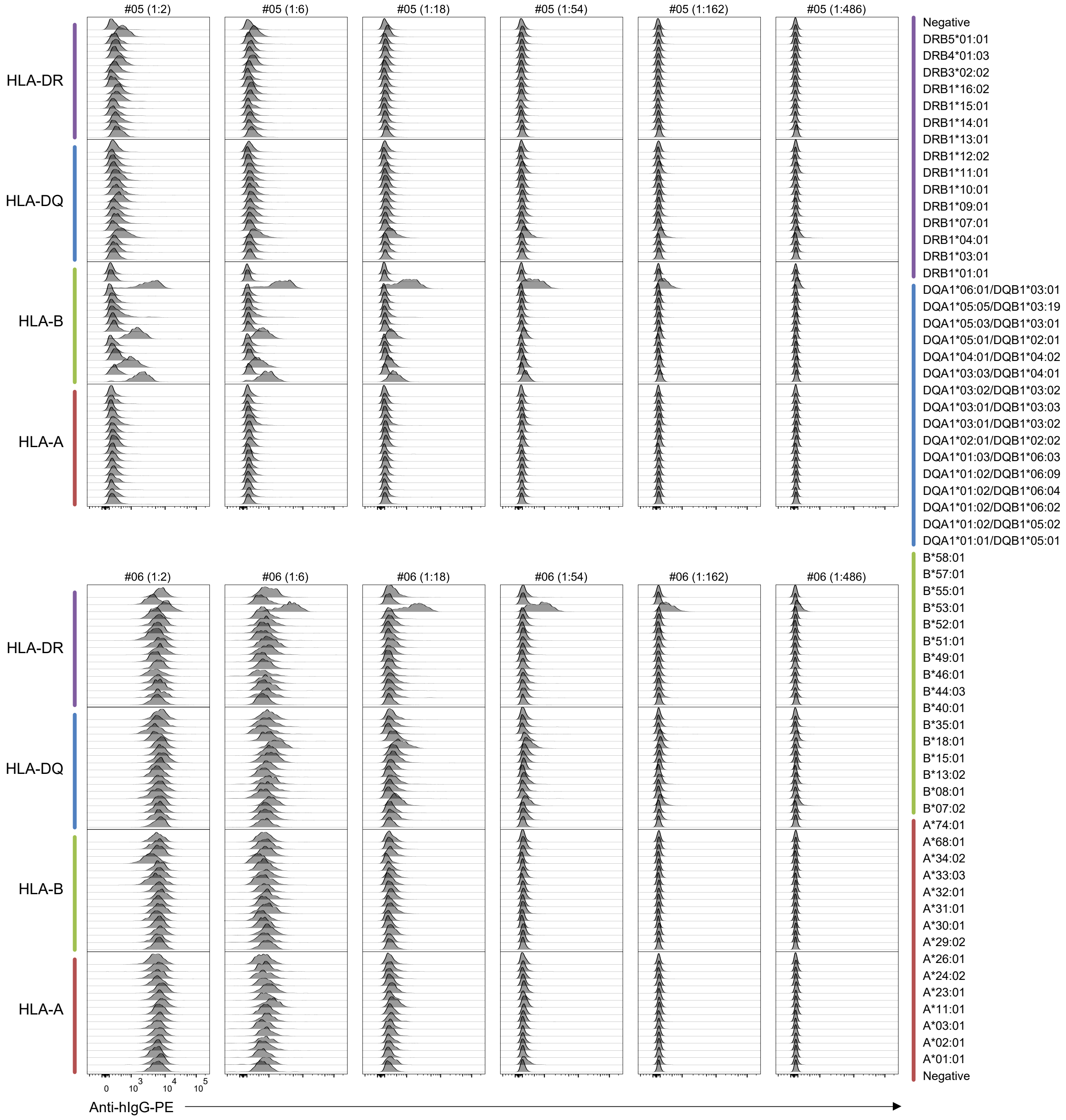

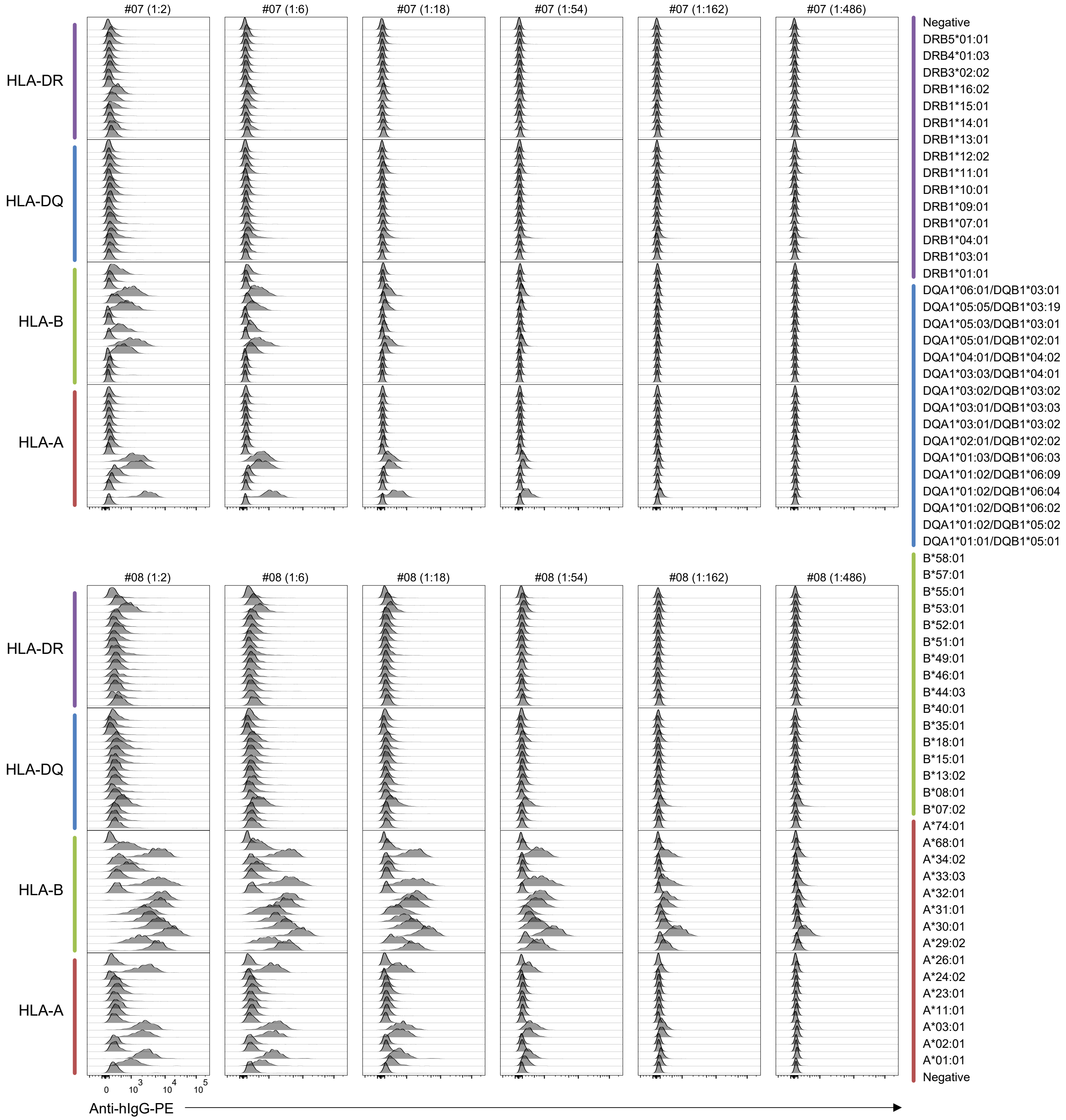

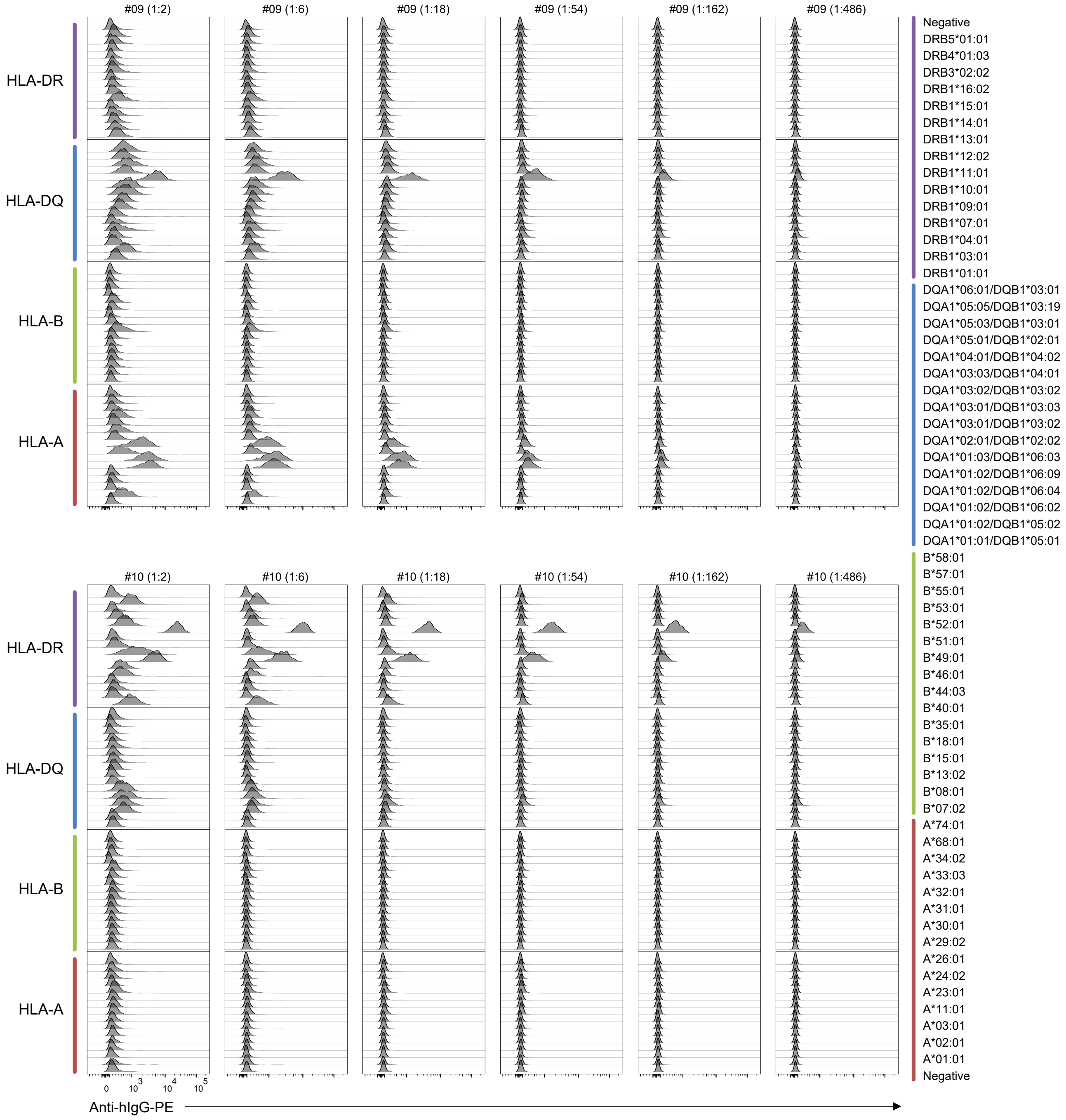

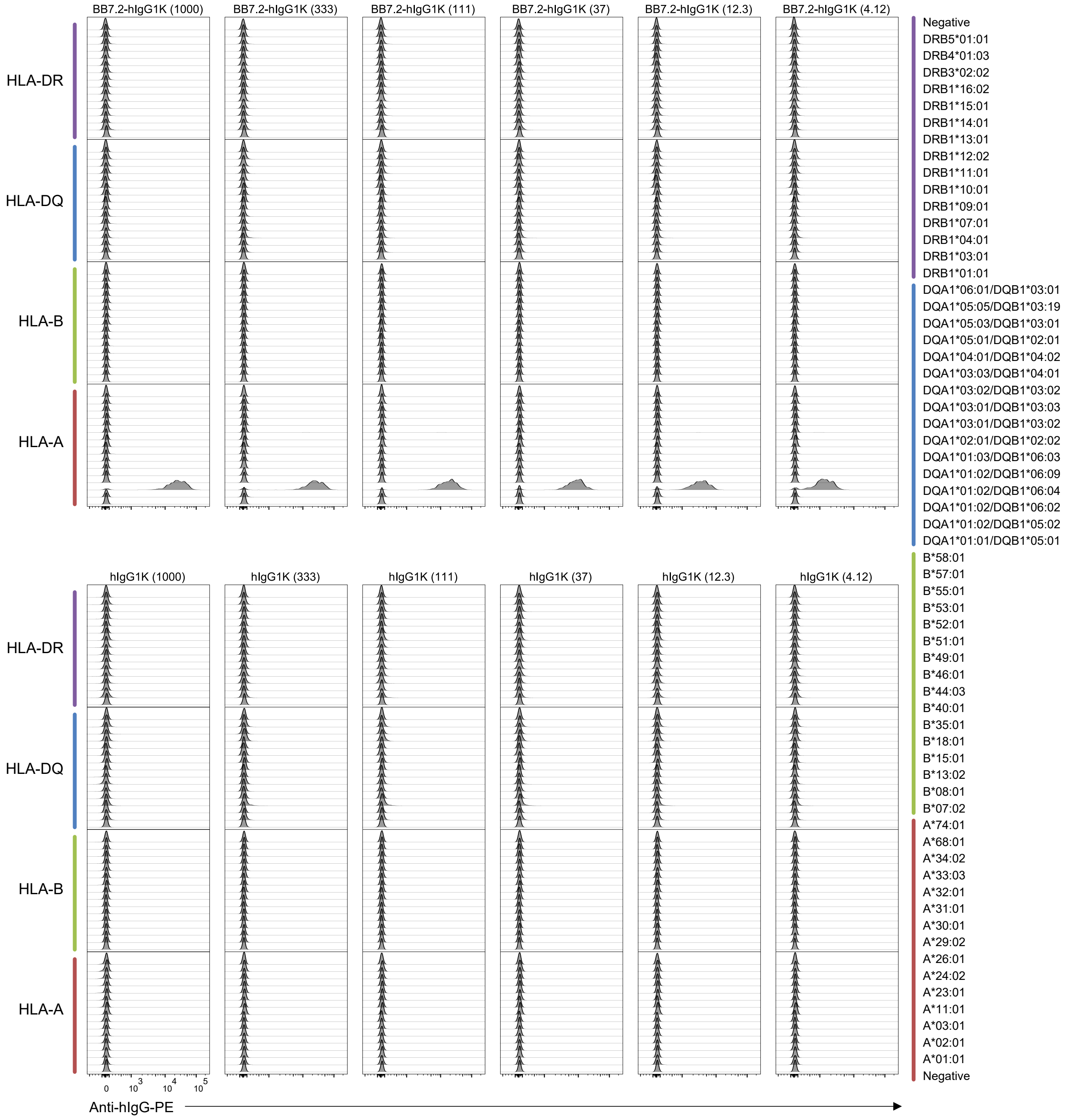

### Supplementary Document 2

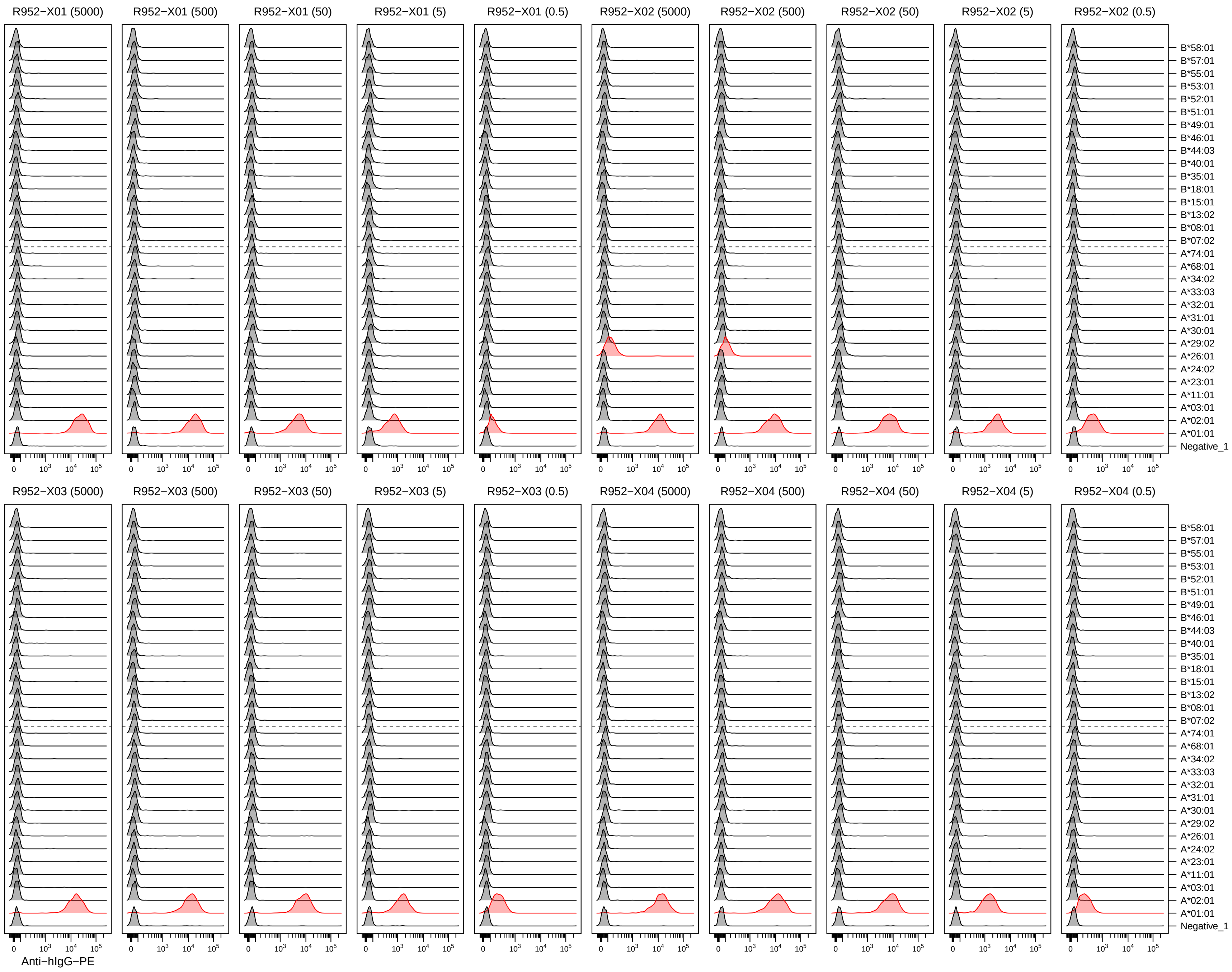

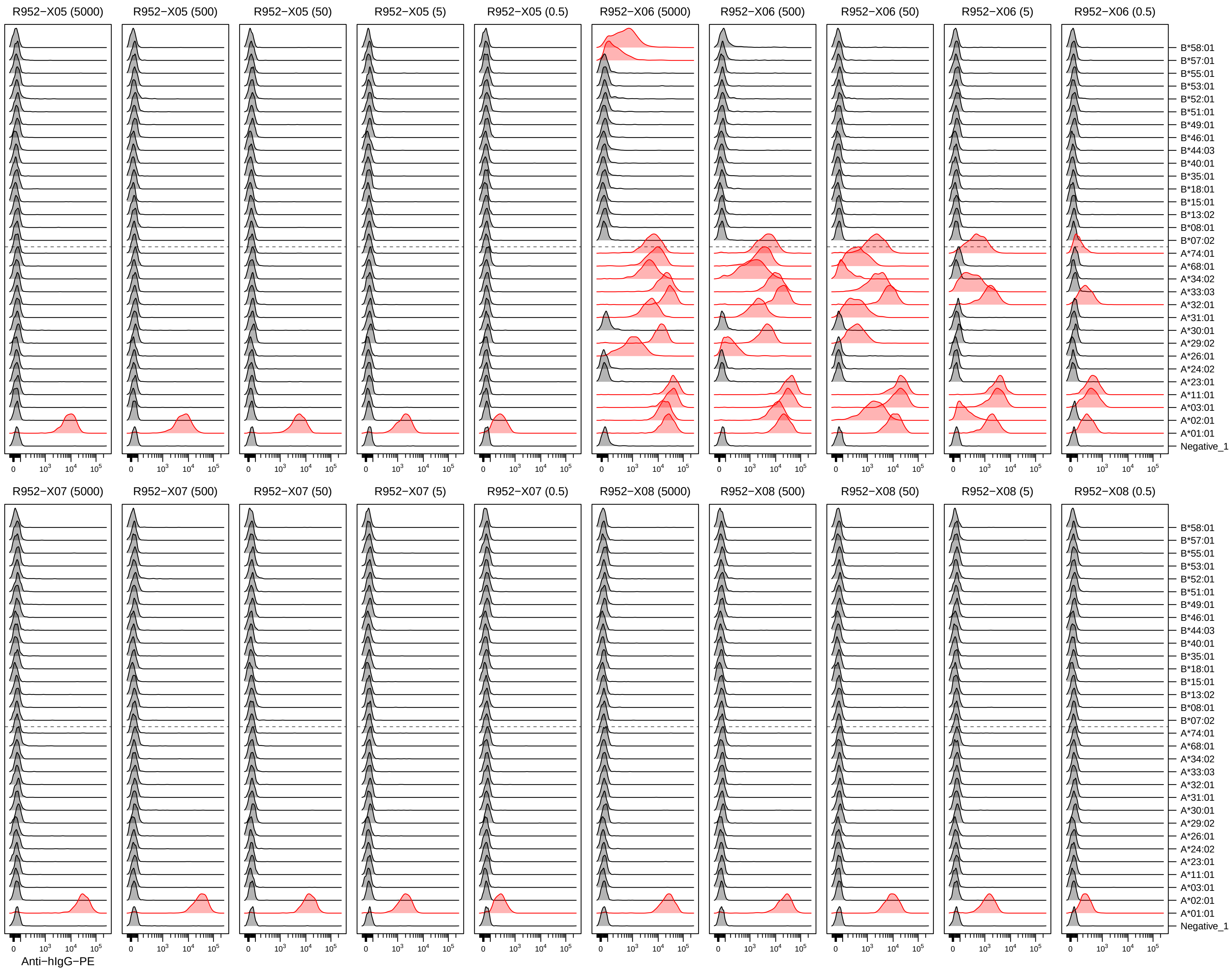

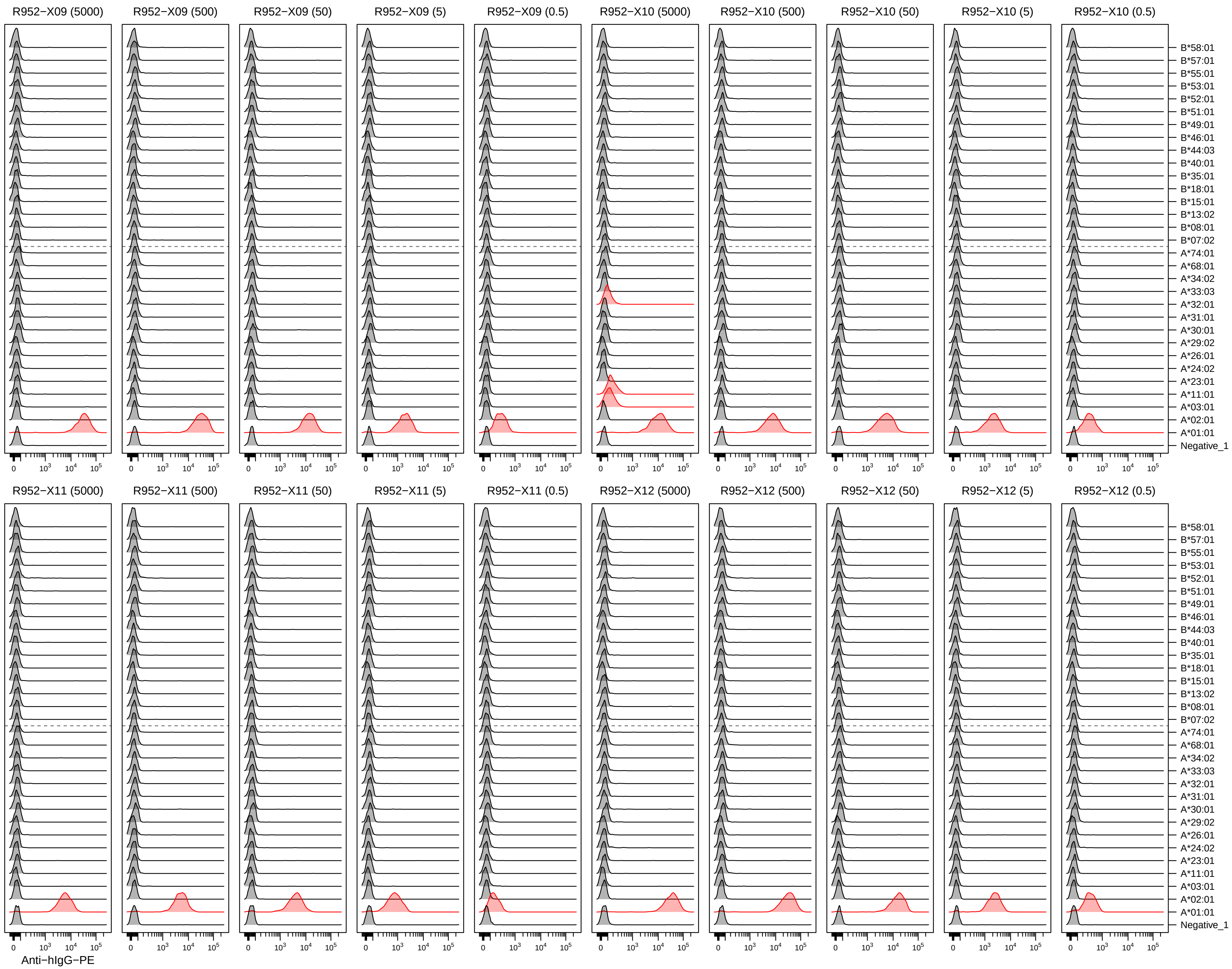

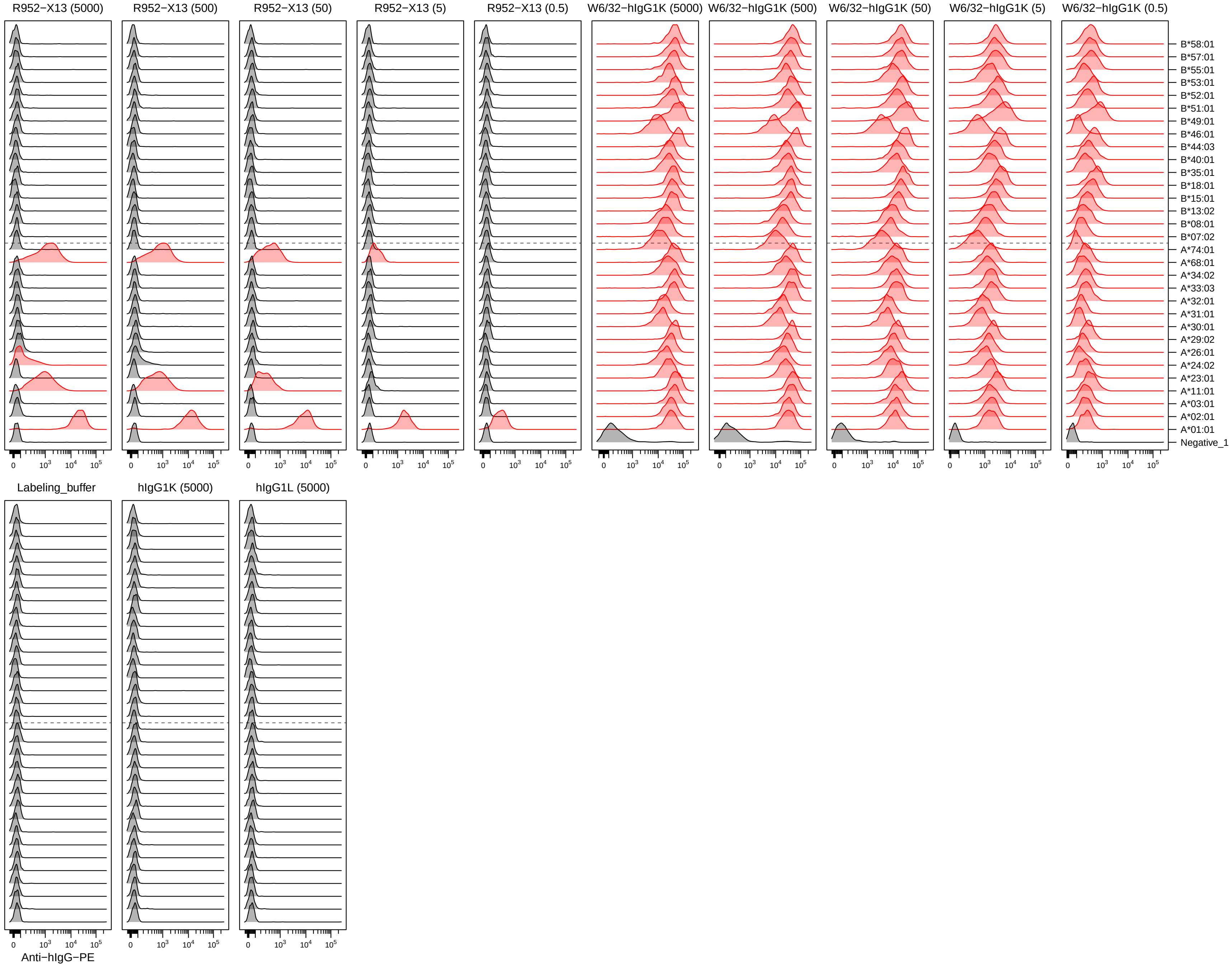
